# FHOD-1/profilin-mediated actin assembly protects sarcomeres against contraction-induced deformation in *C. elegans*

**DOI:** 10.1101/2024.02.29.582848

**Authors:** Michael J. Kimmich, Sumana Sundaramurthy, Meaghan A. Geary, Leila Lesanpezeshki, Curtis V. Yingling, Siva A. Vanapalli, Ryan S. Littlefield, David Pruyne

## Abstract

Formin HOmology Domain 2-containing (FHOD) proteins are a subfamily of actin-organizing formins important for striated muscle development in many animals. We showed previously that absence of the sole FHOD protein, FHOD-1, from *C. elegans* results in thin body-wall muscles with misshapen dense bodies that serve as sarcomere Z-lines. We demonstrate here that actin polymerization by FHOD-1 is required for its function in muscle development, and that FHOD-1 cooperates with profilin PFN-3 for dense body morphogenesis, and profilins PFN-2 and PFN-3 to promote body-wall muscle growth. We further demonstrate dense bodies in *fhod-1* and *pfn-3* mutants are less stable than in wild type animals, having a higher proportion of dynamic protein, and becoming distorted by prolonged muscle contraction. We also observe accumulation of actin depolymerization factor/cofilin homolog UNC-60B in body-wall muscle of these mutants. Such accumulations may indicate targeted disassembly of thin filaments dislodged from unstable dense bodies, and may account for the abnormally slow growth and reduced strength of body-wall muscle in *fhod-1* mutants. Overall, these results show the importance of FHOD protein-mediated actin assembly to forming stable sarcomere Z-lines, and identify profilin as a new contributor to FHOD activity in striated muscle development.

## INTRODUCTION

Striated muscle across the animal kingdom is defined by regularly repeating arrangements of well-ordered contractile units called sarcomeres, whose presence allows for rapid muscle contraction. The ends of sarcomeres, called Z-lines, serve as sites where actin-based thin filaments are crosslinked and anchored in place, while the free ends of thin filaments project toward the sarcomere center to interdigitate with myosin-based thick filaments (Henderson et al., 2017). During contraction, thick filaments pull the thin filaments, sliding the two filament types past each other and shortening the sarcomere (Huxley & Hanson, 1954).

Formin HOmology-2 Domain-containing (FHOD) proteins are a subgroup of actin-organizing formin proteins that have been implicated in striated muscle development in animals ranging from humans to nematodes. However, the contribution of FHOD homologs to sarcomere organization varies from organism to organism. Among mammals, FHOD3 is essential for sarcomere organization in cardiac muscle. Cultured cardiomyocytes derived from rat or from human induced pluripotent stem cells are unable to assemble mature sarcomeres when FHOD3 is knocked down (Taniguchi et al., 2009; Iskratsch et al., 2010; Fenix et al., 2018). In mice deleted for the *fhod3* gene, cardiomyocytes make immature plasma membrane-associated sarcomeres, but these fail to mature and accumulate much lower levels of filamentous actin (F-actin) than in wild type mice (Kan-O et al., 2012). *Drosophila* indirect flight muscles (IFMs) also strongly depend on the sole fly FHOD protein (called FHOS or FHOD), with FHOS knockout IFMs assembling less F-actin and failing to make sarcomeres (Shwartz et al., 2016). Study of FHOS knockdown later in development in fly IFMs showed FHOD is also required to incorporate new thin filaments into growing sarcomeres, and to elongate the barbed ends of thin filaments already present in sarcomeres (Shwartz et al., 2016). In contrast to these examples, the striated body-wall muscle (BWM) of the simple nematode *Caenorhabditis elegans* is able to assemble sarcomeres even in absence of its sole FHOD protein, FHOD-1. Worms lacking FHOD-1 assemble fewer sarcomeres per muscle cell, but these sarcomeres are functional, and crawling or swimming are not grossly affected (Mi-Mi et al., 2012; Mi-Mi & Pruyne, 2015). The ability to form functional sarcomeres in these worms is not due to redundancy with any of the other five worm formins, as we have shown only the formin CYK-1 also affects BWM development, but through pleiotropic effects on overall body growth, while the remaining formins play no apparent role in BWM development (Mi-Mi et al., 2012; Sundaramurthy et al., 2020).

Formins, including the FHOD proteins, are able to stimulate actin polymerization. Functioning as dimers, formins nucleate new actin filaments from monomers through their formin homolog-2 (FH2) domains, often in conjunction with additional actin-binding sites in the adjacent carboxy-terminal (C-terminal) extension or “tail” (Pruyne et al., 2002; Sagot et al., 2002; Gould et al., 2011; Vizcarra et al., 2014; Bremer et al., 2024). The ring-shaped FH2 dimer remains wrapped around the terminal actin subunits of the growing barbed end (Kovar & Pollard, 2004; Maufront et al., 2023). Depending on the formin, the effects of the FH2 dimer can vary from blocking further actin incorporation (“tight capping”) to allowing actin monomer incorporation at the barbed end without displacement of the formin (“processive capping”) (Aydin et al., 2018). The formin homology-1 (FH1) domain also contributes to actin assembly by recruiting the small actin monomer-binding protein, profilin, through multiple poly-proline motifs, and transferring profilin-bound actin monomers to the FH2-bound barbed end (Kovar et al., 2003; Romero et al., 2004; Moseley et al., 2004). For many formins, including FHOD proteins, nucleation and processive capping depend on a universally conserved isoleucine of the FH2 domain (Xu et al., 2004; Patel et al., 2018; Antoku et al., 2019). For mammalian FHOD3, mutation of this isoleucine (I1127) to alanine prevents the formin from supporting sarcomere formation in cultured cardiomyocytes (Taniguchi et al., 2009), while overexpression of FHOD3(I1127A) in the mouse heart recapitulates *fhod3* gene knockout phenotypes (Fujimoto et al., 2016). Similarly, *Drosophila* FHOS(I966A) is only partially functional, organizing actin filaments into small sarcomeres, but failing to support further sarcomere thickening or elongation (Shwartz et al., 2016). Thus, mammalian FHOD3 and *Drosophila* FHOS require their actin assembly activity in order to function properly in their respective muscles. Considering worm FHOD-1 is not required to assemble functional sarcomeres in BWM, we questioned whether worm FHOD-1 might function independently of actin assembly activity.

Here, we demonstrate that in fact actin polymerization by FHOD-1 is critical to proper BWM development in *C. elegans*. Moreover, we show that FHOD-1 works in conjunction with profilin, the first demonstration of formin/profilin cooperation in a striated muscle model. Finally, we show FHOD-1 and profilin are critical for ensuring the ability of Z-line structures of BWM sarcomeres to resist severe deformation by the forces of muscle contraction.

## RESULTS

### A mutant FHOD-1 predicted to be defective for actin filament assembly does not support BWM growth or normal dense body morphogenesis

The striated BWMs of *C. elegans* are strips of flat muscle cells that stretch from nose to tail. Each muscle cell is tightly adherent to a basement membrane shared with the overlying hypodermis (Waterston, 1988). F-actin- and myosin-rich sarcomeres are arranged in obliquely oriented striations that define the spindle shapes of these cells. Most BWM cells arise during embryogenesis, and grow through larval development and early adulthood, accumulating additional striations over time. We previously demonstrated a requirement for FHOD-1 for normal BWM growth based on the effects of RNA interference (RNAi) or deletion alleles, including *fhod-1(tm2363)* predicted to eliminate part of the FH2 domain and introduce a frameshift that prevents translation of downstream sequence (referred to here as *fhod-1(ΔFH2)*, Fig.1A) (Mi-Mi et al., 2012). In *fhod-1(RNAi)* or *fhod-1(ΔFH2)* animals, the spindle-shaped muscle cells are narrower than those in wild type animals when measured at their widest point (e.g., Fig.1B, *solid arrows*), which also results in narrower BWMs (e.g., Fig.1B, *dashed arrows*).

**Figure 1.**
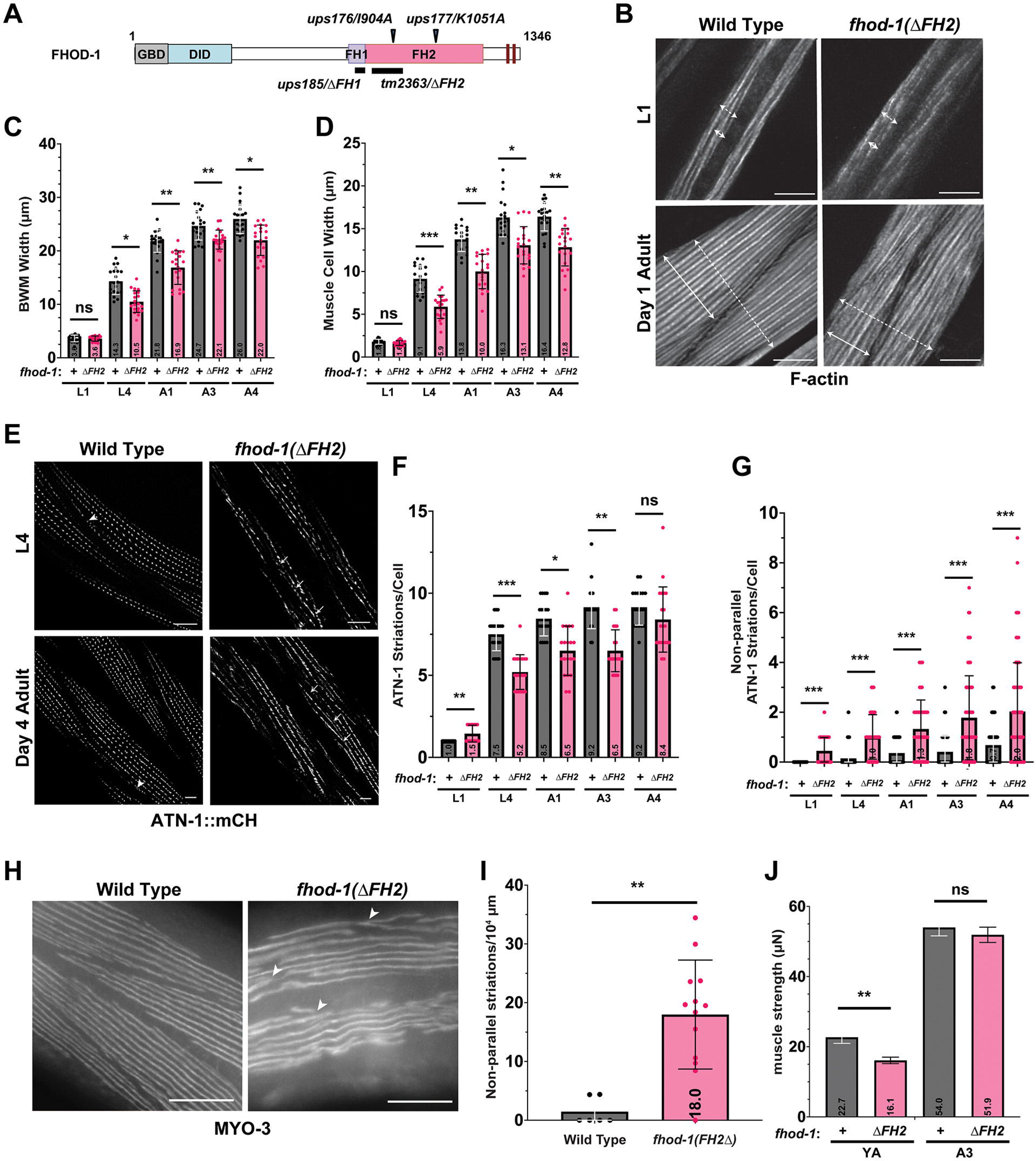
FHOD-1 is required for normal and timely formation of striations during BWM growth. **(A)** Predicted effects of missense mutations (*black arrowheads*) or deletions (*black bars*) in mutants used in this study relative to FHOD-1 structural domains GTPase-binding domain (GBD), diaphanous inhibitory domain (DID), formin homology-1 domain (FH1), formin homology-2 domain (FH2), and two predicted diaphanous autoregulatory domains (*dark red bars*). Numbers indicate amino acid residues of FHOD-1 isoform A. **(B)** Maximum intensity projections of deconvolved confocal z-stacks showing BWMs in dorsal views of phalloidin-stained animals, with examples of the widths of BWMs (*dashed arrows*) and of individual muscle cells (*solid arrows*) indicated. Scale bars, 6 µm. **(C)** BWM widths and **(D)** muscle cell widths were measured in phalloidin-stained worms of larval stages L1 and L4, and animals after one day of adulthood (A1), three days (A3), or four days (A4) (n = 10 animals per genotype per age per trial, 4 BWMs or 4 muscle cells per animal, in two independent trials). Growth of muscle cells in *fhod-1(ΔFH2)* animals lags behind that of wild types. **(E)** Maximum intensity projections of deconvolved confocal z-stacks showing BWMs in dorsal views of animals expressing ATN-1::mCH dense body marker. Intersecting non-parallel striations are more commonly observed in *fhod-1(ΔFH2)* animals (*arrows*) than in wild type animals (*arrowheads*). Scale bars, 6 µm. **(F)** Numbers of ANT-1::mCH-marked striations per muscle cell and **(G)** numbers of non-parallel ANT-1::mCH-marked striations per muscle cell were counted for indicated ages and genotypes (n = 10 animals per genotype per age per trial, 4 muscle cells per animal, in two independent trials). Muscle cells in *fhod-1(ΔFH2)* animals add new striations slower than wild type animals, but accumulate more non-parallel striations. **(H)** Wide-field dorsal views of worms immunostained for MYO-3 show that intersections between non-parallel striations (*arrowheads*) are more prevalent in *fhod-1(ΔFH2)* animals. Scale bars, 50 µm. **(I)** Shown are numbers of non-parallel MYO-3 striations counted per combined measured length of MYO-3 striations visible in wide-field views (n = 6 wild type or 13 *fhod-1(ΔFH2)* worms in one trial). **(J)** Muscle strengths of young (day 0) adults and day 3 adults were measured using the Nemaflex platform. Young adult *fhod-1(ΔFH2)* animals are weaker than wild types, but come close to matching wild type animal strength by the third day of adulthood. Numerical results shown as individual measures and averages ± standard deviation, except **J** showing mean ± SEM. (*) *p* < 0.05; (**) *p* < 0.01; (***) *p* < 0.001; (ns, not significant) *p* > 0.05.

Expanding on our previous analysis (Mi-Mi et al., 2012), we confirmed BWM grows more slowly in larval and adult *fhod-1(ΔFH2)* animals compared to wild type, but observed *fhod-1(ΔFH2)* mutants proportionately narrow their muscle size deficiency in older adults (Fig.1B-D). Examining the Z-line marker α-actinin (ATN-1 in *C. elegans*) to delineate the oblique striations of BWM cells, we observed that this slow growth of muscle cells in *fhod-1(ΔFH2)* animals correlates with slower addition of new striations (Fig.1E,F). Additionally, as compared to wild type animals, more striations formed in *fhod-1(ΔFH2)* animals are misoriented, making fork-like intersections (Fig.1E,G). This defect is also observable when the central A band marker, muscle myosin II heavy chain A/MYO-3, is examined (Fig.1H,I).

While *fhod-1(ΔFH2)* mutants are not grossly deficient at crawling or swimming, we tested whether *fhod-1(ΔFH2)* mutant BWM is quantitatively weaker using the NemaFlex platform, which calculates forces exerted by individual worms based on the degree to which they are able to deflect micropillars (Rahman et al., 2018). Consistent with their ∼ 30% narrower BWMs, young adult *fhod-1(ΔFH2)* animals exert 29% less force, while by the third day of adulthood, *fhod-1(ΔFH2)* BWM is only 10% narrower than wild type, and these animals apply a statistically insignificant 4% less force than wild type (Fig.1J). Thus, thinning of *fhod-1(ΔFH2)* BWM correlates well with muscle weakness, and reflects slower and imperfect addition of new striations during muscle cell growth.

To test the importance of the actin filament assembly activity of FHOD-1 in promoting muscle cell growth, we used CRISPR-Cas9 to introduce FH2 domain missense mutations I904A and K1051A into the endogenous *fhod-1* gene (Fig.1A). Both mutations target universally conserved actin-binding residues, with mutation of the isoleucine to alanine eliminating actin filament assembly activity in many formins, and mutation of the lysine to alanine typically having a milder effect (Xu et al., 2004; Ramabhadran et al., 2012). Immunoblot of whole worm extracts using an antibody raised against the FHOD-1 FH2 domain shows *fhod-1(I904A)* and *fhod-1(K1051A)* worms express FHOD-1 protein at the correct predicted molecular weights (Fig.2A), including a higher molecular weight isoform we have shown previously is expressed in BWM (Refai et al., 2018). In contrast, extracts from *fhod-1(ΔFH2)* animals show only nonspecific immunoreactivity, consistent with our previous observations (Mi-Mi et al., 2012). Staining for F-actin showed that BWM in *fhod-1(I904A)* animals are narrower than wild type, but identical to BWM of age-matched *fhod-1(ΔFH2)* worms, and have nearly identically narrowed muscle cells with fewer striations than wild type (Fig.2B-D). In contrast, BWM of *fhod-1(K1051A)* worms are not significantly different from wild type (Fig.2B-D). Consistent with this, in tests for the ability of worms to burrow through 26% pluronic gel (Lesanpezeshki et al., 2019), *fhod-1(ΔFH2)* and *fhod-1(I904A)* mutants perform worse than wild type animals to a very similar degree, while *fhod-1(K1051A)* worms are nearly as efficient as wild type animals (Fig.2E). Thus, the ability to assemble actin filaments appears to be critical for FHOD-1 function in promoting muscle cell growth and normal BWM performance.

**Figure 2.**
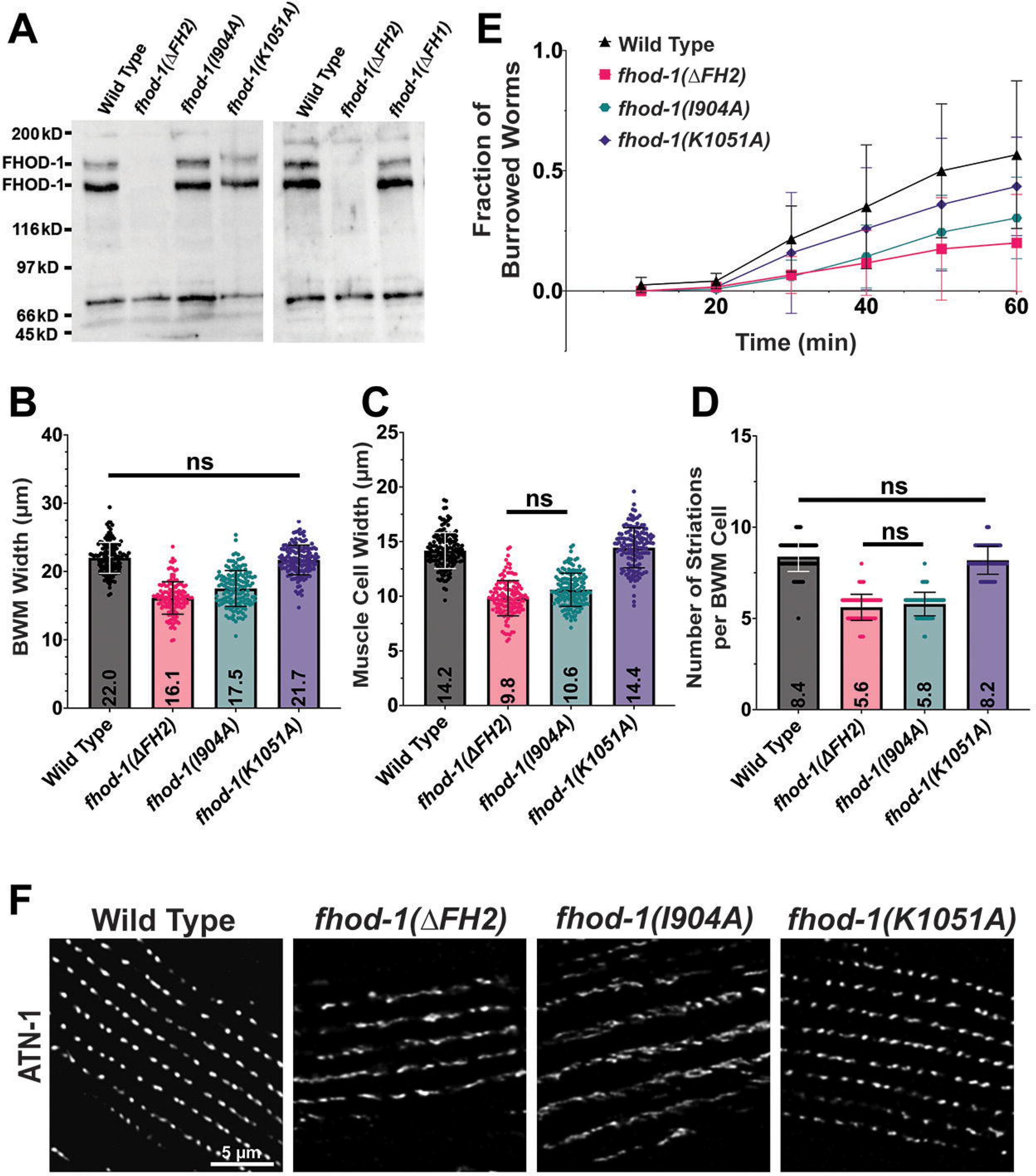
FHOD-1-stimulated actin filament polymerization is required for BWM growth and dense body morphogenesis. **(A)** Two anti-FHOD-1 immunoblots of whole worm extracts show expected ∼150 kDa and ∼165 kDa FHOD-1 isoforms in wild types and in animals bearing missense mutations *fhod-1(I904A)* and *fhod-1(K1051A)* and deletion mutation *fhod-1(ΔFH1)*, but not in animals bearing the deletion and frameshift mutation *fhod-1(ΔFH2)*. Previously observed nonspecific immunoreactivity to lower molecular weight proteins is visible for all strains. **(B)** BWM widths and **(C)** widths of individual muscle cells were measured, and **(D)** numbers of striations per muscle cell were counted in phalloidin-stained day 1 adults (n = 20 animals per genotype per trial, two to four BWMs or muscle cells per animal, in three independent trials). Muscle cells of *fhod-1(I904A)* animals are nearly identically to those of *fhod-1(ΔFH2)* mutants, while *fhod-1(K1051A)* muscle cells are nearly wild type. Numerical results shown as individual measures and averages ± standard deviation. All comparisons are statistically significant, *p* < 0.001, except those indicated (ns, not significant, *p* > 0.05). **(E)** Fraction of day 1 adult worms able to successfully burrow through 26% pluronic gel was determined every 10 min for 1 h (n = 30 worms per genotype per trial, in four independent trials). Results shown as averages ± standard deviation. No differences were statistically significant owing to large trial-to-trial differences, but *fhod-1(I904A)* and *fhod-1(ΔFH2)* animals consistently perform poorly compared to wild type and *fhod-1(K1051A)* animals in each trial. **(F)** Maximum intensity projections of confocal z-stacks show dorsal views of day 1 adults immunostained for dense body marker ATN-1. Dense bodies appear regular in shape and spacing in wild type animals, but irregular in shape and spacing in *fhod-1(ΔFH2)* and *fhod-1(I904A)* mutants. Dense bodies in *fhod-1(K1051A)* animals are very similar to those in wild type animals. Scale bar, 5 µm.

We have also previously demonstrated that *fhod-1(ΔFH2)* animals form misshapen dense bodies (Mi-Mi & Pruyne, 2015; Sundaramurthy et al., 2020). Dense bodies play two roles in BWM, serving as sarcomere Z-lines to anchor thin filaments, and as costameres to anchor sarcomeres to the plasma membrane (Gieseler et al., 2018). When viewed by immunostain for α-actinin/ATN-1, dense bodies are peg shaped structures arranged with semi-regular spacing along striations in wild type animals, whereas dense bodies in adult *fhod-1(ΔFH2)* mutants appear fragmented and irregular in shape, with irregular spacing along striations (Sundaramurthy et al., 2020). Immunostain for ATN-1 showed adult worms bearing *fhod-1(I904A)* have irregular dense bodies identical to those of *fhod-1(ΔFH2)* animals, while dense bodies in *fhod-1(K1051A)* worms appear more regular in shape and spacing, similar to those of wild type animals (Fig.2F). These data suggest FHOD-1 mediated actin assembly is essential for proper dense body morphology, with mutation of I904 having the greatest effect.

### Profilin contributes to BWM cell growth and dense body formation similar to FHOD-1

The *C. elegans* genome encodes three profilins: PFN-1, which is essential and expressed embryonically and in adult neurons, vulva, spermatheca, and myoepithelial sheath of the gonad; PFN-2, which is non-essential and expressed primarily in pharyngeal muscle, intestine, and spermatheca; and PFN-3, which is non-essential and expressed primarily in BWM (Polet et al., 2006). Worms bearing the whole-gene deletion *pfn-3(tm1362)*, referred to here as *pfn-3(Δ)*, had also been previously shown to perform worse in pluronic gel burrowing assays than wild type animals (Lesanpezeshki et al., 2021). Additional, *pfn-3(Δ)* animals were reported to have some neighboring dense bodies that appear to merge with each other, while RNAi against *pfn-2* in the *pfn-3(Δ)* background resulted in sarcomeres that were somewhat wider than normal in 10-20% of treated worms (Polet et al., 2006). To directly compare profilin- and *fhod-1-*deficient BWM, we stained F-actin in worms bearing *pfn-3(Δ)*, as well as worms bearing the whole-gene deletion *pfn-2(ok458)*, referred to here as *pfn-2(Δ)*, or animals deleted for both *pfn-2* and *pfn-3*. BWM of *pfn-2(Δ)* animals is nearly identical to that of age-matched wild type animals, with same-sized muscle cells with the same number of striations per cell, while *pfn-3(Δ)* animals average one fewer striation per muscle cell compared to wild type, and double *pfn-2(Δ) pfn-3(Δ)* mutants have modestly narrower muscle cells and an average of two fewer striations per cell (Fig.3A-F). These effects are similar to those *fhod-1(ΔFH2)*, but of lesser magnitude. ATN-1 immunostain revealed that dense bodies in *pfn-2(Δ)* animals are normal, but that dense bodies are somewhat irregular in shape and spacing in *pfn-3(Δ)* mutants and in double *pfn-2(Δ) pfn-3(Δ)* mutants, although often not as irregular in appearance as in *fhod-1(ΔFH2)* animals (Fig.3G). Thus, PFN-3 and to a lesser extent PFN-2 contribute to BWM growth, and PFN-3 contributes to proper dense body morphogenesis, similar to FHOD-1.

**Figure 3.**
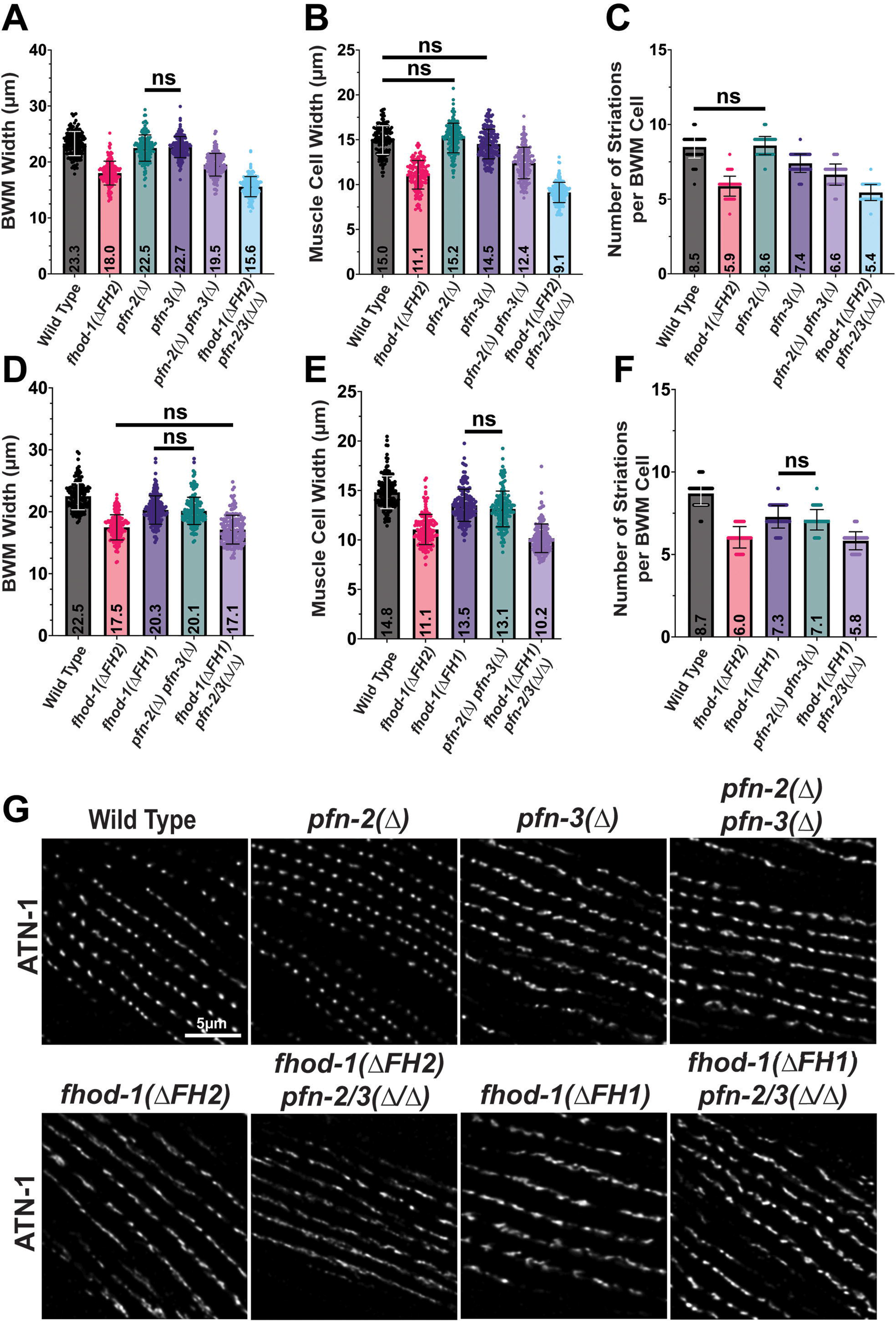
Profilins and FHOD-1 FH1 domain are required for normal BWM cell growth and dense body morphogenesis. **(A-C)** Comparison of single and double profilin mutants to *fhod-1(ΔFH2)* mutants for **(A)** BWM width, **(B)** widths of individual muscle cells, and **(C)** numbers of striations per muscle cell, in day 1 adults (n = 20 animals per genotype per trial, two to four BWMs or muscle cells per animal, in three independent trials). Loss of the profilin genes *pfn-2* and *pfn-3* contribute to all three BWM metrics, although their combined effects are less than that of *fhod-1(ΔFH2)*. **(D-F)** Comparison of *fhod-1(ΔFH1)* mutants to single and double profilin mutants for **(D)** BWM width, **(E)** widths of individual muscle cells, and **(F)** numbers of striations per muscle cell, in day 1 adults (n = 20 animals per genotype per trial, two to four BWMs or muscle cells per animal, in three independent trials). The effects of the loss of profilin genes *pfn-2* and *pfn-3* on BWM metrics closely match the effects of *fhod-1(ΔFH1)*. Results shown as individual measures and averages ± standard deviation. All comparisons are statistically significant, *p* < 0.001, except those indicated (ns, not significant, *p* > 0.05). **(G)** Maximum intensity projections of confocal z-stacks show dorsal views of day 1 adults immunostained for dense body marker ATN-1. Dense bodies appear similarly irregular in *pfn-3(Δ)* mutants, double *pfn-2(Δ) pfn-3(Δ)* mutants, and *fhod-1(ΔFH1)* mutants. Scale bar, 5 µm.

If profilin cooperates with FHOD-1 to promote actin filament assembly for BWM development, we would expect the FHOD-1 FH1 domain to also be important for BWM development. To test this, we used CRISPR-Cas9 to delete from the endogenous *fhod-1* gene the three poly-proline stretches in the FH1 domain predicted to bind profilin (Fig.1A). Immunoblotting verified *fhod-1(ΔFH1)* animals produce FHOD-1 protein of the expected molecular weights (Fig. 2A). By F-actin stain, BWM in *fhod-1(ΔFH1)* animals is narrower than that of wild type animals, with identical muscle cell widths and number of striations per muscle cell as age-matched double *pfn-2(Δ) pfn-3(Δ)* mutants (Fig.3D-F). By immunostain for ATN-1, dense bodies in *fhod-1(FH1Δ)* animals are highly similar to those of *pfn-3(Δ)* mutants, being somewhat irregular is shape and spacing (Fig.4E). Thus, eliminating the predicted profilin-binding sites in the FHOD-1 FH1 domain has nearly identical effects on BWM development as elimination of PFN-2 and PFN-3.

**Figure 4.**
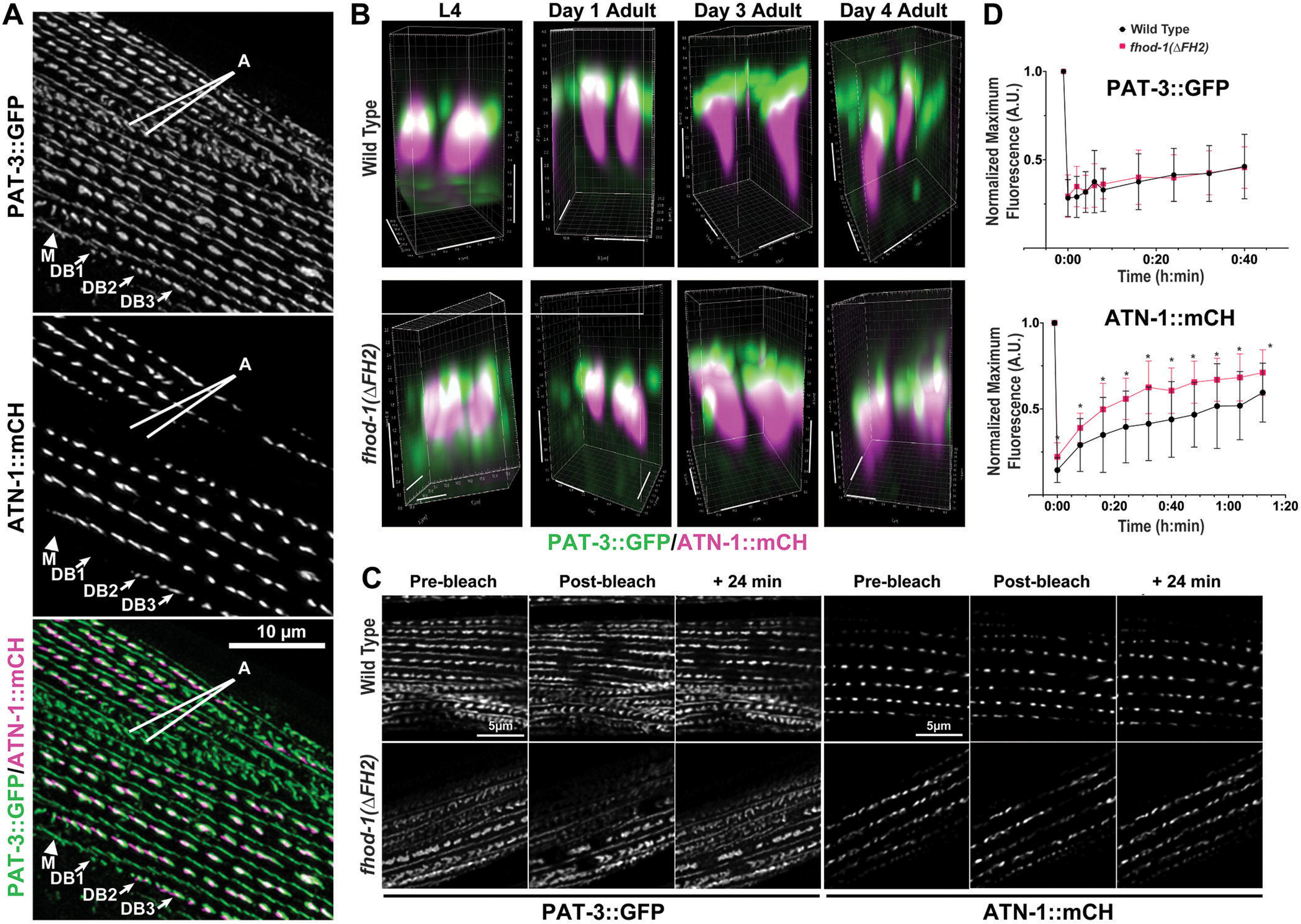
α-actinin is more mobile in dense bodies formed in the absence of FHOD-1. **(A)** Maximum intensity projection of a deconvolved confocal z-stack of a day 1 adult expressing PAT-3::GFP and ATN-1::mCH. Both markers are associated with dense bodies, although dense body composition varies along striations, starting as small dense bodies at striation ends (DB1) with PAT-3::GFP only, and becoming progressively larger (DB2 to DB3) and associated with ATN-1::mCH further from striation ends. PAT-3::GFP is also associated with M-lines (M) and attachment plaques (A), both of which can be identified based on their differing morphologies. Scale bar, 10 µm. **(B)** Three-dimensional reconstructions from deconvolved confocal z-stacks show dense bodies from PAT-3::GFP/ATN-1::mCH-expressing worms. PAT-3::GFP is present at the plasma membrane and ATN-1::mCH is present in elongated projections embedded in the sarcomere lattice. Dense bodies tend to be shorter and wider in *fhod-1(ΔFH2)* animals. Scale bars, 1 µm. **(C)** Single confocal images of live L4 larvae show PAT-3::GFP- or ATN-1::mCH-marked dense bodies prior to photobleaching, immediately after photobleaching targeted dense bodies, and after 24 min recovery. Scale bars, 5 µm. **(D)** Quantification of FRAP of dense bodies imaged as in **C** (for PAT-3::GFP, n = 4 worms of each genotype, 4 dense bodies per animal; for ATN-1::mCH, n = 5 worms of each genotype, 4 dense bodies per animal). PAT-3::GFP recovers slowly and to similar degrees in wild type and *fhod-1(ΔFH2)* animals, while ATN-1::mCH recovers more rapidly in both strains, but to a greater degree in *fhod-1(ΔFH2)* animals. Results shown are averages ± standard deviation. (*) *p* < 0.05. No differences were significant between wild type and *fhod-1(ΔFH2)* animals at any time point for PAT-3::GFP (*p* > 0.05).

To test whether profilins and FHOD-1 contribute to muscle development through common or separate mechanisms, we examined BWM of animals mutated simultaneously for *fhod-1*, *pfn-2*, and *pfn-3*. We expected that if profilins contribute to BWM development strictly through FHOD-1-mediated actin filament assembly, the effect of these combined mutations would be no worse that the effect of mutating *fhod-1* alone. Conversely, if profilins contribute to BWM development through additional FHOD-1-independent mechanisms, then the phenotype of triple mutants would exceed that of mutating *fhod-1* alone. Based on immunostain for ATN-1, dense bodies in triple *fhod-1(ΔFH1) pfn-2(Δ) pfn-3(Δ)* mutants appear similar to those of *fhod-1(ΔFH1)* or *pfn-3(Δ)* mutants (Fig.3G). Similarly, dense bodies in triple *fhod-1(ΔFH2) pfn-2(Δ) pfn-3(Δ)* appear similar to those of *fhod-1(ΔFH2)* mutants. Based on this, profilin, and particularly PFN-3, likely contributes to dense body morphogenesis through interaction with the FHOD-1 FH1 domain. However, the effects of triple mutants on BWM growth were less straightforward. F-actin stain showed muscle cells of triple *fhod-1(ΔFH1) pfn-2(Δ) pfn-3(Δ)* mutants are narrower and have fewer striations than those of *fhod-1(FH1Δ)* or of double *pfn-2(Δ) pfn-3(Δ)* mutants (Fig.3D-F). Similarly, BWM growth was more reduced in triple *fhod-1(ΔFH2) pfn-2(Δ) pfn-3(Δ)* mutants than in *fhod-1(ΔFH2)* mutants (Fig.3A-C). These results suggest the profilins PFN-2 and PFN-3 contribute to BWM growth independently of FHOD-1, although they may also contribute to BWM growth through FHOD-1-mediated actin filament assembly.

### Dense bodies formed in the absence of PFN-3 or FHOD-1 are unstable

*C. elegans* dense bodies have a layered organization similar to focal adhesions, with a membrane-proximal zone containing α-integrin/PAT-2 and β-integrin/PAT-3, an intermediate layer rich in vinculin/DEB-1, and an elongated segment rich in α-actinin/ATN-1 for thin filament attachment (Francis & Waterston, 1985; Barstead & Waterston, 1989; Gettner et al., 1995; Moulder et al., 2010). To better understand the effects of FHOD-1 on dense body dynamics, we examined animals expressing functional GFP-tagged β-integrin (GFP::PAT-3) and functional mCherry-tagged α-actinin (ATN-1::mCH) (Fig.4A,B, S1). As described previously, GFP::PAT-3 decorates dense bodies as well as M-lines and muscle cell-to-muscle cell adhesions called attachment plaques (Plenefisch et al., 2000). ATN-1::mCH also decorates most dense bodies, but is absent from smaller dense bodies found at the ends of striations at the edges of muscle cells (Fig.4A), a trait observed in wild type and *fhod-1(ΔFH2)* animals. Extending our observations of misshapen dense bodies in adult animals (Fig.2F, 3G), ATN-1::mCH-positive dense bodies of *fhod-1(ΔFH2)* of all ages are more lobulated and variably shaped than in wild type worms (Figs 4B, S1). Past analysis of dense body spacing in ATN-1-immunostained animals by fast Fourier transform (FFT) had shown dense bodies in wild type young adults exhibit a semi-regular 1-µm spacing along striations, while *fhod-1(ΔFH2)* young adults have no preferred spacing (Sundaramurthy et al., 2020). Consistent with and expanding upon this, FFT analysis of dense bodies in ATN-1::mCH-expressing larvae and adults of varying ages showed wild type animals exhibit regular dense body spacing throughout growth, trending from ∼ 0.7 µm between dense bodies in the youngest stage-1 larvae (L1) to > 1.8 µm at the fourth day of adulthood (Table 1, Fig.S2). Conversely, FFT analysis of *fhod-1(ΔFH2)* dense bodies shows no preferred spacing through larval development, and only weak and inconsistent preferred spacing in older *fhod-1(ΔFH2)* adults (Table 1, Fig.S2). Thus, dense body malformation is present in *fhod-1(ΔFH2)* animals throughout development.

**Table 1.** Dense body spacing along striations.

| Age | Predominant Dense Body Spacing<sup>†</sup> |  |
| --- | --- | --- |
| | Wild Type | *fhod-1*($\Delta FH2$) |
| L1 | 0.73 $\mu\text{m}$, n/a<sup>‡</sup> | n/a<sup>‡</sup>, n/a<sup>‡</sup> |
| L4 | 1.17 $\mu\text{m}$, 1.45 $\mu\text{m}$ | n/a<sup>‡</sup>, n/a<sup>‡</sup> |
| Day 1 Adult | 1.50 $\mu\text{m}$, 2.38 $\mu\text{m}$ | n/a<sup>‡</sup>, 2.00 $\mu\text{m}$ |
| Day 3 Adult | 1.58 $\mu\text{m}$, 2.09 $\mu\text{m}$ | 3.74 $\mu\text{m}$, n/a<sup>‡</sup> |
| Day 4 Adult | 1.84 $\mu\text{m}$, 2.25 $\mu\text{m}$ | 2.64 $\mu\text{m}$, n/a<sup>‡</sup> |

<sup>†</sup> Values shown are calculated as the inverse of the primary frequency ( $\mu\text{m}^{-1}$ ) determined by FFT analysis of ATN-1::mCH fluorescence along striations (Fig.S2). Values from each of two trials are shown, determined from fluorescence profiles of all dense body-containing striations of two muscle cells (for L1 larvae) or eight dense body-containing striations in one muscle cell (for other ages) in each of ten animals per strain per developmental stage.
<sup>‡</sup> No primary frequency was present, or the calculated predominant dense body spacing spanned multiple dense bodies.

To test whether protein dynamics in *fhod-1(ΔFH2)* dense bodies are abnormal, we performed fluorescence recovery after photobleaching (FRAP). Individual dense bodies in immobilized L4 larvae were photobleached and monitored for recovery for 40 or 72 min (Fig.4C,D). GFP::PAT-3 showed a slow, linear recovery of fluorescence that did not saturate, and did not differ between wild type and *fhod-1(ΔFH2)* animals (Fig.4C,D). ATN-1::mCH recovered more rapidly in both strains, with *fhod-1(ΔFH2)* dense bodies showing a greater degree of fluorescence recovery (Fig.4C,D), suggesting the α-actinin-rich portion of dense bodies that anchors thin filaments is more dynamic in absence of FHOD-1. We reasoned that if dense bodies were defective in anchoring thin filaments in *fhod-1(ΔFH2)* animals, we might observe enhanced recruitment of the muscle-specific actin depolymerization factor (ADF)/cofilin homolog, UNC-60B, a protein essential for actin filament turnover in BWM (Ono et al., 1999). In wild type animals, we observe UNC-60B in low amounts along the Z-lines (Fig.5), as shown previously (Ono et al., 1999). However, in *fhod-1(ΔFH2)* animals, UNC-60B also accumulates at the ends of the spindle-shaped muscle cells of BWM (Fig.5, *arrows*). We had not previously noted F-actin accumulation at muscle cell ends in *fhod-1(ΔFH2)* animals, but ADF/cofilin and phalloidin bind competitively to actin filaments (Nishida et al., 1987; Ono et al., 1996; Ono & Ono, 2009). Thus, we probed for actin by immunostain, and observed UNC-60B at the *fhod-1(ΔFH2)* muscle cell ends overlaps with actin (Fig.5, *arrows*). We observed similar but fainter UNC-60B accumulation BWM in double *pfn-2(Δ) pfn-3(Δ)* mutants, with faint overlapping actin immunostain (Fig.5). Distinct from *fhod-1(ΔFH2)* animals, profilin mutants also accumulate UNC-60B puncta in the muscle cell cytoplasm beneath the layer of sarcomeres (Fig.5, *arrowheads in Organellar Layer*), consistent with profilins playing at least some FHOD-1-independent roles in BWM development.

**Figure 5.**
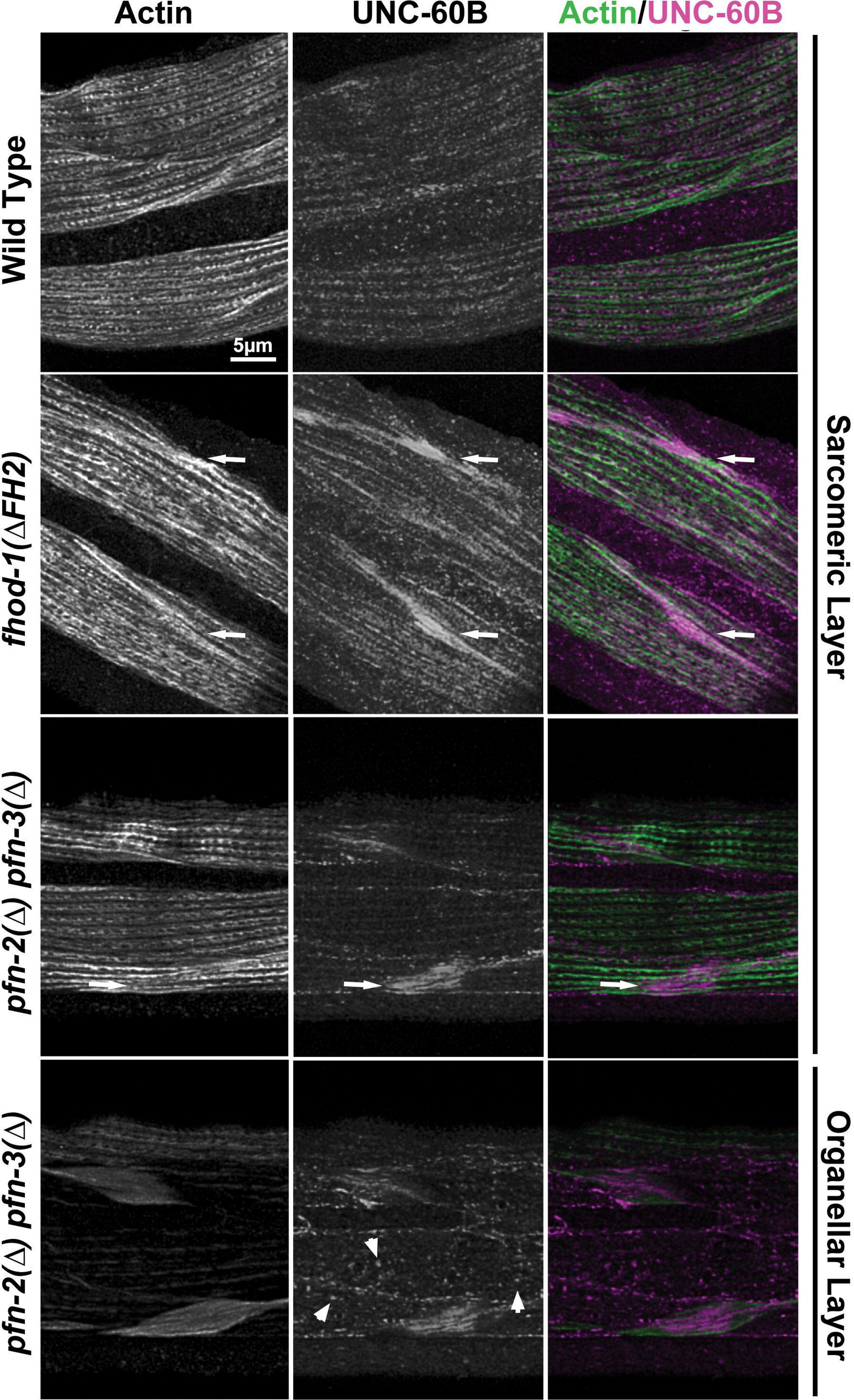
Muscle cells lacking FHOD-1 or PFN-3 accumulate ADF/cofilin-associated actin filaments. Maximum intensity projections of deconvolved confocal z-stacks show day 1 adults immunostained for actin and UNC-60B. UNC-60B and actin are organized into striations in BWM. In *fhod-1(ΔFH2)* animals and, to a lesser extent, in *pfn-2(Δ) pfn-3(Δ)* animals, UNC-60B also accumulates at the narrow ends of muscle cells (*arrows*). Additionally, *pfn-2(Δ) pfn-3(Δ)* animals accumulate small puncta of UNC-60B (*arrowheads*) in the portion of the muscle cell cytoplasm occupied by organelles, below the layer of sarcomeres. Scale bar, 5 µm.

Based on our evidence that the thin filament-binding portion of dense bodies in *fhod-1(ΔFH2)* mutants is more dynamic than in wild type, we hypothesized that the deformed shape of *fhod-1(ΔFH2)* dense bodies might arise from an inability to resist pulling forces during muscle contraction. To test this, worms were grown in the presence of 2.5 mM levamisole, a nicotinic acetylcholine receptor agonist. This maintained BWM in a partially contracted state, as verified by a tauter appearance of the animals, although the worms were still able to crawl. After 24 h treatment, we observed 100% wild type and 95% *fhod-1(ΔFH2)* levamisole-treated worms remained alive. Where sarcomeres in wild type BWM are largely unaffected by this treatment, sarcomeres of levamisole-treated *fhod-1(ΔFH2)* animals are highly distorted compared to untreated controls (Fig.6A). Dense bodies in treated *fhod-1(ΔFH2)* animals, as viewed through ATN-1::mCH and PAT-3::GFP, are stretched to form nearly continuous-appearing Z-lines rather than discontinuous bodies (Fig.6A, *arrows*), and striations are contorted rather than straight. There is also an increase in diffusely PAT-3::GFP across the muscle cell membrane in levamisole-treated *fhod-1(ΔFH2)* BWM, while M-line-associated PAT-3::GFP is reduced (Fig.6A). We also tested whether prolonged muscle relaxation would reverse dense body morphological defects in *fhod-1(ΔFH2)* worms grown in the presence of 10 mM muscimol, a γ-aminobutyric acid receptor agonist. However, after 24 h, this treatment did not dramatically improve *fhod-1(ΔFH2)* dense bodies, suggesting dense bodies deformation may be irreversible (Fig.6A).

**Figure 6.**
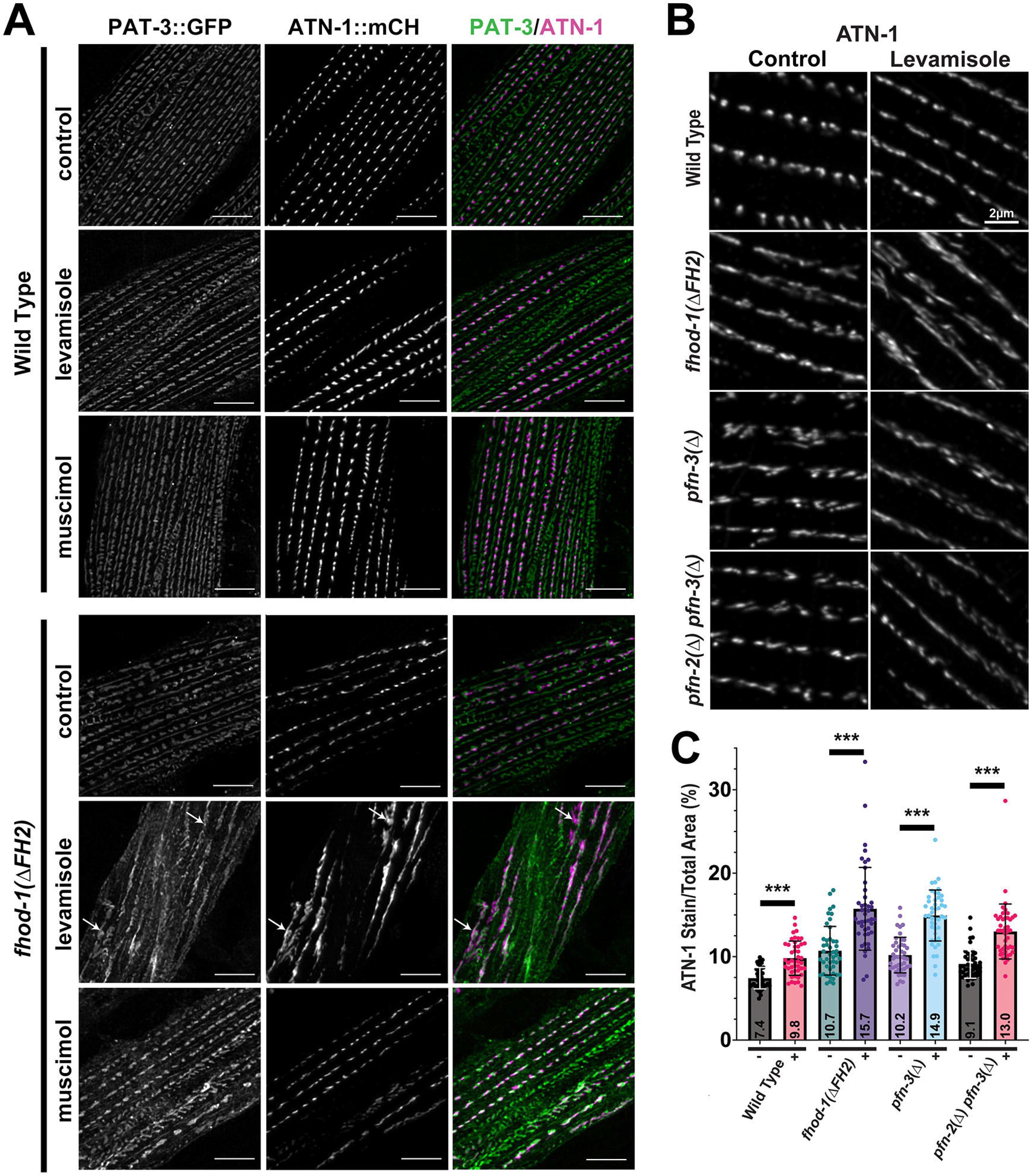
Dense bodies formed in the absence of FHOD-1 or PFN-2/PFN-3 are highly deformed by prolonged contraction. **(A)** Maximum intensity projections of deconvolved confocal z-stacks show day 1 adults expressing PAT-3::GFP and ATN-1::mCH. Dense bodies in wild type animals are only mildly distorted after 24 h treatment with 2.5 mM levamisole, which stimulates muscle contraction, or with 10 µM muscimol, which inhibits muscle contraction. In contrast, dense bodies in *fhod-1(ΔFH2)* animals are highly distorted by levamisole treatment, forming nearly continuous stripes of dense body material (*arrows*). Scale bars, 6 µm. **(B)** Maximum intensity projections of deconvolved confocal z-stacks show day 1 adults immunostained for ATN-1 after 24 h treatment with or without 2.5 mM levamisole. Scale bar, 2 µm. **(C)** The effects of levamisole treatment were measured from images as in **B** by determining the total area occupied by ATN-1 in maximum intensity projections of untreated (-) and levamisole-treated (+) animals (n = 1 image from each of 15 worms per genotype per condition per trial, with three independent trials). All strains are significantly changed by levamisole treatment (*p* < 0.001), although levamisole treatment more strongly distorts dense bodies in *fhod-1(ΔFH2)*, *pfn-3(Δ)*, and double *pfn-2(Δ) pfn-3(Δ)* mutants than in wild type animals.

To quantify the effects of levamisole treatment on dense body morphology, we immunostained control- or levamisole-treated worms for ATN-1, and measured the percent surface area taken up by the dense body marker in two-dimensional projections of sarcomeres (Fig.6B,C). By this metric, wild type animals exhibit very modest dense body deformation that results in ATN-1 occupying 2.4% more of the total area after levamisole treatment. Dense body deformation was quantitatively greater in *fhod-1(ΔFH2)* mutants, with ATN-1 occupying 5% more of the total area (Fig.6B,C). Similarly, dense bodies levamisole-treated in *pfn-3(Δ)* mutants occupy 4.7% greater area than in untreated animals, and those in levamisole-treated double *pfn-2(Δ) pfn-3(Δ)* occupy 3.9% greater area (Fig.6B,C), supporting a common FHOD-1/profilin pathway for dense body morphogenesis.

Considering FHOD-1 is important for the ability of dense bodies to resist prolonged muscle contraction, we tested whether FHOD-1 is actively recruited to dense bodies during levamisole treatment. Under normal conditions, a functional GFP-tagged FHOD-1 (FHOD-1::GFP) localizes to small bodies along the lateral edges of BWM cells (Fig.7, *white arrows*), and fainter bodies within sarcomeres, particularly around and between dense bodies (Fig.7, *cyan arrows*). This localization is consistent with that of endogenous FHOD-1 (Mi-Mi et al., 2012). Localized FHOD-1::GFP remains visible through larval development and young adulthood, but fades to nearly undetectable levels by the third day of adulthood, when BWM growth has ceased (Fig.7, *L4* versus *Day 3 Adult*). Interestingly, in levamisole-treated wild type animals, FHOD-1::GFP is not localized in BWM sarcomeres, unlike untreated control animals (Fig.7, *control* versus *levamisole*). This is consistent with our observation that muscle cells of levamisole-treated animals are strongly inhibited for adding growth (Fig.S3). Together, these results suggest that FHOD-1 does not strengthen dense bodies by being actively recruited during prolonged contraction. Instead, we suggest dense body stability is established in a FHOD-1-dependent manner during dense body formation, when FHOD-1 is present in and near sarcomeres.

**Figure 7.**
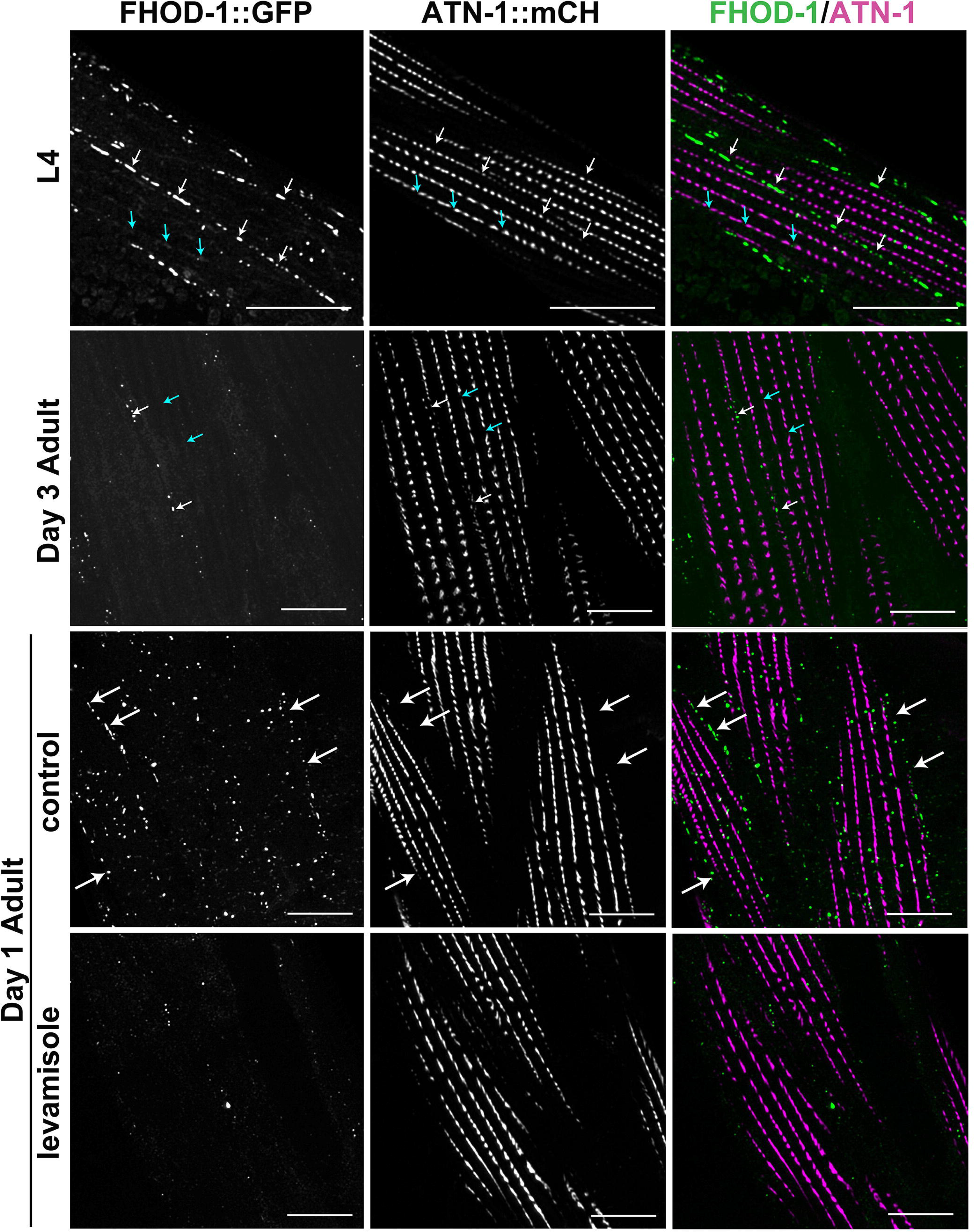
FHOD-1 is present among sarcomeres during BWM growth, but not under conditions of prolonged contraction. Maximum intensity projections of deconvolved confocal z-stacks show dorsal views of animals expressing FHOD-1::GFP and ATN-1::mCH. In L4 larvae, FHOD-1 is present in bodies along the edges of BWM cells (*white arrows*) that are distinct from dense bodies, and in faint striations (*cyan arrows*) that intersect dense bodies. By day 3 adulthood, most localized FHOD-1::GFP is gone. Treatment of day 1 adults with levamisole for 24 h results in delocalization of FHOD-1::GFP, as compared to untreated controls. Scale bars, 10 µm.

## DISCUSSION

### *In vivo* cooperation between FHOD-1 and profilin in striated muscle development

We had shown previously that among six members of the formin family of actin organizing proteins in *C. elegans*, only FHOD-1 directly contributes to the development of the body-wall muscle (BWM) in a cell autonomous manner (Mi-Mi et al., 2012; Sundaramurthy et al., 2020). In absence of FHOD-1, muscle cells of BWM add new striations more slowly during development, resulting in slower BWM growth (Fig.1E,F). Additionally, dense bodies, which serve as sarcomere Z-lines, are misshapen and irregular in shape and spacing when formed in the absence of FHOD-1 (Figs 4A, S1). However, unlike mouse and fly models for FHOD-dependent muscle development, sarcomeres of FHOD-1-deficient BWMs are functional, and contain abundant actin-based thin filaments (Mi-Mi & Pruyne, 2015). This led to the question of whether FHOD-1 functions independently of the ability to stimulate the actin filament polymerization. We show here that a *fhod-1* mutant bearing the FH2 domain missense mutation I904A, predicted disrupt actin filament assembly, phenocopies a functionally null *fhod-1* allele for muscle cell growth, dense body morphology, and BWM performance in burrowing assays (Fig.2). A second FH2 missense mutation K1051A, predicted to have milder impact on actin filament assembly, had little observable effect on these phenotypes.

Many formins have been shown to work with the actin monomer-binding protein profilin to promote actin filament assembly *in vitro*, using the FH1 domain to recruit profilin with bound actin to the growing ends of elongating actin filaments (Coutemanche, 2018). We provide here the first *in vivo* evidence of cooperation between a FHOD protein and profilin in striated muscle development. Absence of the profilin PFN-3, which is primarily expressed in BWM, results in dense bodies that are irregular in shape and spacing (Fig.2F, 3G) (Polet et al., 2006). In direct comparisons, these dense body defects appear very similar to those in *fhod-1* mutants, and particularly in animals disrupted for the FHOD-1 FH1 domain predicted to bind profilin (Fig.3G). Moreover, we saw no additive effects on dense bodies with simultaneous loss of PFN-3 and the FHOD-1 FH1 domain, suggesting PFN-3 contributes to dense body morphogenesis through its interaction with the FHOD-1 FH1 domain (Fig.3G).

We also observed both profilins PFN-2 and PFN-3 contribute redundantly to BWM growth. Thus, worms lacking PFN-3 have only very modest BWM growth defects, but worms lacking both PFN-2 and PFN-3 have greater defects (Fig.3A-C). However, combined absence of PFN-2, PFN-3, and FHOD-1 results in BWM growth defects even greater than those caused by absence of profilins or FHOD-1 alone, indicating the profilins must make FHOD-1-independent contributions to BWM growth (Fig.3A-C). Considering profilins also mediate ATP/ADP-nucleotide exchange for actin, suppress pointed end elongation of actin filaments, and interact with a variety of other proteins (Pimm et al., 2020), this result is not surprising. Perhaps related to this, we noted that worms lacking both PFN-2 and PFN-3 accumulate small puncta of the ADF/cofilin homolog UNC-60B in their cytoplasm (Fig.5), possibly indicating a wider disturbance in cytoskeletal organization in the muscle cell. Additionally, PFN-2 is predominantly expressed in pharyngeal muscle and intestine (Polet et al., 2006), such that some effects of profilin loss on BWM growth might be secondary to overall reduced nutrition and growth. This is consistent with our observation that worms triply mutated for *fhod-1*, *pfn-2*, and *pfn-3* are smaller than age-matched single or double mutant counterparts.

Although cooperation between FHOD proteins and profilin has not been demonstrated in other striated muscle systems, we believe this is likely to be a conserved mechanism. Supporting this, normal sarcomere growth in indirect flight muscles (IFMs) of *Drosophila* requires that its FHOD protein, FHOS, be competent to assemble actin filaments (Shwartz et al., 2016), and FHOS cooperates with profilin *in vitro* to promote actin filament assembly (Patel et al., 2018). There is indirect evidence for cooperation between mammalian FHOD3 and profilin-1 in cultured rat cardiomyocyte, with studies showing that either hyperactivation of FHOD3 or overexpression of profilin-1 is sufficient to induce hypertrophy (Kooij et al., 2016; Zhou et al., 2017). Additionally, in humans, autosomal dominant mutations in *fhod3* cause between 1-2% of inherited cases of hypertrophic cardiomyopathy (HCM) (Ochoa et al., 2018), and expression of FHOD3 is generally elevated in HCM hearts in humans (Wooten et al., 2013), while mouse models of HCM show elevated profilin-1 expression (Kooij et al., 2016). We suggest FHOD3 and profilin likely function as partners in promoting mammalian cardiomyocyte hypertrophy, analogous to how FHOD-1 and profilin promote worm BWM growth.

### FHOD-1 and profilin impart stability onto dense bodies

When viewed by electron microscopy, dense bodies in wild type worms appear as discrete electron-dense structures, but in *fhod-1(ΔFH2)* mutants appear partially fragmented (Mi-Mi & Pruyne, 2015). Considering thin filaments are anchored to dense bodies, we hypothesized their partial fragmentation in *fhod-1(ΔFH2)* animals might result from dense bodies being less physically resilient, and unable to resist pulling forces on thin filaments during muscle contraction. We show here by several metrics that dense bodies are less stable in worms lacking functional FHOD-1 or PFN-3, and this particularly affects the α-actinin-rich portion of dense bodies where thin filaments are anchored. Confirming that muscle contraction is a deforming force, dense bodies in BWM of *fhod-1* and *pfn-3* mutants become highly distorted during prolonged pharmacologically induced contraction (Fig.6). Even under normal conditions, a higher proportion of α-actinin in dense bodies is exchangeable with the cytoplasm in *fhod-1(ΔFH2)* mutant muscle cells (Fig.4C,D), suggesting less of the substance of the dense body is firmly anchored in place. Possibly related, *fhod-1* and *pfn-3* mutant muscle cells accumulate ADF/cofilin with actin at their cell ends (Fig.5). An intriguing hypothesis is that this represents accumulations of thin filaments that have been dislodged from dense bodies and targeted for ADF/cofilin-mediated depolymerization.

Based on FHOD-1 localization in BWM, we suggest FHOD-1 imparts resiliency onto dense bodies as the dense bodies form, rather than acting continuously or in response to acute increases in pulling forces on dense bodies. In growing BWM, FHOD-1 localizes to small bodies along muscle cell edges (Fig.7, *white arrows*) (Mi-Mi et al., 2012). Interestingly, we observe that dense bodies near muscle cell edges, but not in the muscle cell interior, are frequently small and lack ATN-1 (Fig.4A), suggesting they may be immature. FHOD-1 does not colocalize with these dense bodies, but its proximity at the muscle cell edge might permit FHOD-1-generated actin filaments to affect nascent dense bodies as they form. Extant dense bodies in BWM continue to grow in size through larval development and early adulthood (Figs 4B, S1) (Moerman & Williams, 2006). FHOD-1 is also present in small amounts around and between these dense bodies (Fig.7, *cyan arrows* in *L4*) (Mi-Mi et al., 2012), potentially allowing FHOD-1 to support their on-going growth. However, we argue it is unlikely FHOD-1 actively maintains dense body integrity after dense bodies have finished assembling, as FHOD-1 largely no longer localizes near dense bodies after BWM growth ceases (Fig.7, *L4* versus *Day 3 Adult*). Additionally, FHOD-1 is not recruited to dense bodies during prolonged muscle contraction, when structural integrity is most challenged (Fig.7). The localization of PFN-3 through these conditions is less clear. Previous immunostain has shown PFN-3 concentrates near the tips of dense bodies in adult BWM (Polet et al., 2006), but its distribution during larval development when BWM growth is greatest remains to be determined. Also to be determined is precisely how FHOD-1/PFN-3-mediated actin filament assembly might strengthen dense bodies. One model is that FHOD-1 and PFN-3 elongate thin filament barbed ends through the full mass of the dense body to ensure they are firmly embedded. In absence of FHOD-1 or PFN-3, thin filaments might only partially penetrate the dense body. As such weakly-anchored thin filaments are pulled, they might tear the outer layers of the dense body away, contributing to dense body deformation and to loss of thin filament anchorage. Alternatively, based on the presence of FHOD-1 between dense bodies, FHOD-1/PFN-3 might polymerize as-yet unidentified nonsarcomeric actin filaments that provide structural stability by linking adjacent dense bodies to each other, analogous to the role played by intermediate filaments that bridge Z-discs in vertebrate striated muscles (Henderson et al., 2017).

We suggest that the narrower shape of muscle cells of *fhod-1* and *pfn-3* mutants are secondary consequences of their dense body fragility. As sarcomeres form and enlarge during BWM growth, subsets of thin filaments in *fhod-1* and *pfn-3* mutants become dislodged and are disassembled. However, a net positive balance of assembly over disassembly allows for formation of new striations at a slower rate, with resulting narrow muscle (Figs 1J, 2E). A base rate of sarcomere assembly failure might explain our previous observation that muscle myosin II heavy chain/MYO-3 is subject to elevated levels of proteasome-driven proteolysis in *fhod-1(ΔFH2)* BWM (Yingling & Pruyne, 2021). With all other factors remaining equal, we would expect muscle strength to be proportional to cross-sectional area (Moss et al., 1997). Despite structural defects, BWM in *fhod-1(ΔFH2)* adults appears to follow this rule. That is, muscle cells in *fhod-1(ΔFH2)* BWM have the same length as in wild type animals, and the same radial thickness when viewed in cross sections (Mi-Mi et al., 2012). Thus, the degree of lateral narrowing in BWM (measured as “width”) correlates directly with changes in cross-sectional area. Strikingly, the measured muscle strength observed in *fhod-1(ΔFH2)* animals correlates very well with the width of their BWM relative to wild type animals at several ages (Fig.1J), suggesting their BWM weakness as young animals is also a secondary consequence of slowed BWM growth.

In several ways, the relationship between FHOD-1 and dense bodies resembles the relationship between FHOD3 and sarcomeres in the mouse heart. During heart development in the mouse, cardiomyocytes assemble immature sarcomeres in stress fiber-like premyofibrils that support an early heartbeat (Kan-O et al., 2012). Normally, premyofibrils mature by forming proper sarcomere Z-discs and incorporating additional thin filaments, but in *fhod3* knockout cardiomyocytes, sarcomeric organization is lost, and α-actinin- and F-actin-rich aggregates form (Kan-O et al., 2012). An intriguing possibility is that in the absence of FHOD3, nascent cardiomyocyte sarcomeres are unstable, similar to *fhod-1* mutant dense bodies. As the mouse heart begins beating when premyofibrils are forming in the mouse heart, it is possible that the growing sarcomeres in the *fhod3* knockout become disrupted by the increasing contractile strength of the developing heart. As a test of this, it would be interesting to determine whether modest reductions in contractility might support more advanced sarcomere maturation in *fhod3-* deficient cardiomyocytes. Another parallel between worm FHOD-1 and mouse FHOD3 is that neither seems strictly required to maintain sarcomeres after they have assembled. Thus, after conditional knockout of FHOD3 from the adult mouse heart, sarcomeres are maintained and the heart becomes mildly enlarged (Ushijima et al., 2018). These commonalities may indicate that despite the difference in the severity of their phenotypes, loss of these conserved proteins in worm and mouse may have very similar mechanistic consequences on sarcomeric organization.

## MATERIALS AND METHODS

### Worms strains and growth conditions

Worms were grown on nematode growth medium (NGM) agar with OP50-1 *Escherichia coli* bacterial food at 20°C following standard protocols (Brenner, 1974). Where needed, worms were age-synchronized by dissolving gravid adults in 1:2 ratio of reagent grade bleach to 5 M NaOH to liberate embryos (Bartel, 1991), and hatching embryos into L1 larvae overnight at 20°C in M9 buffer (Ausubel et al., 2002). To induce prolonged partial muscle contraction or relaxation (for Figs 6,7), OP50-1 culture was mixed with 2.5 mM levamisole, 10 mM muscimol, or no drug (control or vehicle) prior to seeding 500 µl to 60-mm NGM plates, or 250 µl to 35-mm NGM plates, similar to as described (Brouilly et al., 2018). L4 stage larvae were washed once with M9 buffer and added to dried plates for 24 h treatment at 20°C.

All worm strains used in this study are listed in Table 2. Worm strain DWP294 (genotype *rhIs2[pat-3::HA::gfp]*) was previously created (Plenefisch et al., 2000) but was not previously named. CRISPR-Cas9-mediated mutagenesis was performed on worms by injecting the syncytial gonad with Cas9 protein (Integrated DNA Technologies, IDT, Coralville, IA), tracrRNA (IDT), gene-specific crRNAs (IDT), and single-stranded oligodeoxynucleotides (ssODNs) (IDT) for homology-directed repair, as described (Paix et al., 2015). Allele *fhod-1(ups176)*, encoding amino acid change I904A and a novel *MwoI* cut site for PCR-based genotyping, was generated using crRNA [UUU AGU UAG ACC AAU GUU GAG UUU UAG AGC UAU GCU] and ssODN [AAA CTC TAT CTG TTC TTC CTC TGA AAA GAT CGC AAG CAA TCA ACG CAG GTC TAA CTA AAT TGC CAC CGA TCA ACG TCA TCC CTG CAG CAA TTA T]. Allele *fhod-1(ups177)*, encoding K1051A and eliminating an endogenous *BamHI* site, was generated with crRNA [AAC AAA AGC AUC AGA AGU AAG UUU UAG AGC UAU GCU] and ssODN [GGA ACT GAT ATT AAG GGT TTC TAT CTG GAT TAT TTA ACA AAA GCA TCA GAA GTA GCA GAT CCA GTC TAC AAG CAT ACT TTG ACA TAT CAC]. Allele *fhod-1(ups185)*, encoding a deletion of G768 to G805 of the FH1 domain, was generated with crRNAs [CGU GGA GGU CCU GGU GUG UUU UAG AGC UAU GCU] and [AUU CCU CCA CCU CCU CCU CCG UUU UAG AGC UAU GCU], and ssODN [CGG AGA ATG GAA TGC GTG GAG GTC CTG TTG GTG TTA ATT TGC TTA TGA ACG GTA TAA ATC GAG GAG ATA T]. All CRISPR-generated *fhod-1* alleles were verified by sequencing the modified loci, and were outcrossed six times to a clean N2 (wild type) background. All amino acid residue numbers are relative to FHOD-1 isoform A, as currently defined on WormBase http://www.wormbase.org/db/get?name=CE41309#06--10;class=protein.

**Table 2.** Worm strains used in this study.

| Strain | Genotype | Source<sup>†</sup> | Figure(s) |
| --- | --- | --- | --- |
| DWP3 | *qals8001[fhod-1::GFP mini-unc-119(+)]* | Mi-Mi et al., 2012 | Parental stain |
| DWP229 | *atn-1(ftw35) V; rhIs2[pat-3::HA::GFP]* | RSL62, DWP294 | 1E-G, 4A-D, 6A, S1, S2, S3 |
| DWP230 | *fhod-1(tm2363) I; atn-1(ftw35) V; rhIs2[pat-3::HA::GFP]* | XA8001, DWP229 | 1E-G, 4B-D, 6A, S1, S2, S3 |
| DWP231 | *atn-1(ftw35 atn-1::mCH::ICR::GFPnls) V; rhIs2[pat-3::HA::GFP]; qals[fhod-1::GFP mini-unc-119(+)]* | DWP3, RSL62 | 7 |
| DWP284 | *pfn-3(tm1362) X* | N2 outcross of ON167 | 3A-C,G, 6B,C |
| DWP285 | *fhod-1(ups175 [I904A]) I* | CRISPR N2 | 2A-F |
| DWP286 | *fhod-1(ups180 [K1051A]) I* | CRISPR N2 | 2A-F |
| DWP287 | *pfn-2(ok458) X* | N2 outcross of RB694 | 3A-C,G |
| DWP292 | *pfn-3(tm1362) pfn-2(ok458) X* | DWP284, DWP287 | 3A-G, 6B,C |
| DWP294 | *rhIs2[pat-3::HA::GFP]* | Plenefisch et al., 2000 | Parental strain |
| DWP296 | *fhod-1(tm2363) I; pfn-3(tm1362) pfn-2(ok458) X* | XA8001, DWP292 | 3A-C,G |
| DWP303 | *fhod-1(ups185 [ΔFH1]) I* | CRISPR N2 | 3D-G |
| DWP314 | *fhod-1(ups185) I; pfn-3(tm1362) pfn-2(ok458) X* | DWP292, DWP303 | 3D-G |
| N2 | Bristol wild type | CGC<sup>‡</sup> | 1B-D, H-J, 2A-F, 3A-G, 5, 6B,C |
| ON167 | *pfn-3(tm1362) X* | CGC<sup>‡</sup> | Parental strain |
| RB694 | *pfn-2(ok458) X* | CGC<sup>‡</sup> | Parental strain |
| RSL62 | *atn-1(ftw35 [atn-1::mCH::ICR::GFPnls]) V* | Ryan Littlefield | Parental strain |
| XA8001 | *fhod-1(tm2363 [ΔFH2]) I* | Mi-Mi et al., 2012 | 1B-D, H-J, 2A-F, 3A-G, 5, 6B, C |

<sup>†</sup>Reference for initial isolation of strain, or parental strains for strains produced in this study.
<sup>‡</sup>*Caenorhabditis* Genetic Center (University of Minnesota, Minneapolis, MN)

For visualization of α-actinin in live animals, RSL62 was generated by CRISPR-Cas9 mediated homology directed repair (Dickenson et al., 2013) using a co-conversion strategy (Arribere et al., 2014). Briefly, the repair plasmid was generated in two steps: blunt-end cloning of N2 genomic DNA to isolate upstream and downstream homology arms, followed by isothermal assembly to incorporate the bicistronic tagging cassette consisting of *linker-mCherry::ICR::GFP-nls* in-frame at the 3’ end of the *atn-1* coding sequence. The trans-splicing ICR was derived from operon CEOP1032. The *GFP* and *mCherry* sequences were respectively derived from pJA245 and pJA281, which were gifts from Julie Ahringer (Addgene plasmids #21506 and #22513) (Zeiser et al., 2011). The Cas9 sgRNA plasmid was generated according to Arribere and colleagues (2014) using pRB1017, a gift from Andrew Fire (Addgene plasmid #59936). For further details on targeting of *atn-1* with Cas9, see Supplemental Methods. The RSL62 strain was outcrossed five times with wildtype N2 worms without any noticeable change in fluorescence.

### Western blot analysis

Worms were collected and lysed after washing from plates, as previously described (Yingling & Pruyne, 2021). Sample loads for western blot analysis were normalized based on Coomassie brilliant blue stain of extracts after SDS-PAGE. Normalized extract samples were resolved by SDS-PAGE followed by transfer to nitrocellulose for probing with polyclonal antibody DPMSP2 (affinity-purified anti-FHOD-1 FH2 domain; Mi-Mi et al., 2012) diluted 1:200 in 1% milk/Tris-buffered saline, pH 8.3 (Fig.2A).

### Fixation and staining for fluorescence microscopy

Fixation and staining for F-actin with Alexa Fluor 568-phalloidin were as previously (Mi-Mi et al., 2012). Fixation and immunostaining were as previously (Finney & Ruvkun, 1990), but with omission of spermidine-HCl and an increased methanol concentration (75%) during fixation. Monoclonal antibody MH35 (anti-ATN-1) developed by R.H. Waterston (Francis & Waterston, 1985) was a generous gift from P. Hoppe (Western Michigan University, Kalamazoo, MI). Monoclonal antibody 5-6-s (anti-MYO-3) developed by H.F. Epstein (Miller et al., 1983) was obtained through the Developmental Studies Hybridoma Bank, created by the NICHD of the NIH and maintained at The University of Iowa, Department of Biology, Iowa City, IA 52242. Polyclonal anti-UNC-60B (Ono & Ono, 2002) was a gift from S. Ono (Emory University, Atlanta, GA). Monoclonal antibody clone C4 (anti-actin) is commercially available (Thermo Fisher Scientific, Waltham, MA). Primary antibody dilutions were MH35 1:10^4^, 5-6-s 1:10^3^, anti-UNC-70B 1:200, DPMSP2 1:200, and C4 1:50. Secondary antibodies Texas red-conjugated goat anti-rabbit and fluorescein isothiocyanate-conjugated goat-anti-mouse (Rockland Immunochemicals, Limerick, PA) were diluted 1:500.

### Fluorescence microscopy and image analysis

Wide-field fluorescence images (Fig.1H) were acquired using an Eclipse 90i upright microscope (Nikon, Tokyo, Japan) with a CFI Plan Apochromat 40X/NA 1.0 oil immersion objective or CFI Plan Apochromat violet corrected 60X/NA 1.4 oil immersion objective with a Cool-SNAP HA2 digital monochrome charge-coupled device camera (Photometrics, Tucson, AZ) at room temperature, driven by NIS-Elements AR acquisition and analysis software (version 3.1; Nikon). Confocal images (all other figure images) were acquired using an SP8 Laser Scanning Confocal Microscope (Leica, Wetzlar, Germany) driven by LAS X Software (version 3.5.2, build 4758; Leica), and using an HCX Plan Apochromat 63X/NA 1.4 oil immersion lambda objective. Confocal z-stacks of BWMs were collected at 0.1 µm z-intervals (Figs 1B,E, 4A,B, 6A, 7) or 0.3 µm z-intervals (Figs 2F, 3G, 5, 6B) before deconvolution using Huygens Essential software (Huygens compute engine 18.10.0, Scientific Volume Imaging, Hilversum, Netherlands), Classic Maximum Likelihood Estimation deconvolution algorithm, with 40 iterations and a signal/noise ratio of 20. Maximum intensity projections were generated from deconvolved confocal z-stacks using LAS X software or ImageJ (version 2.0.0-rc-65/1.51 g) (Schneider et al., 2012). Images were linearly processed to enhance contrast and false colored in Adobe Photoshop CS4 or CC2018 (Adobe, San Jose, CA). Deconvolved confocal z-stacks were also used to construct 3D renderings (Figs 4B, S1) using Imaris x64 software (version 9.2.1; Bitplane AG, Belfast, United Kingdom).

Measurements of BWM widths and muscle cell widths, and counting of striations per muscle cell were all performed on muscle cells positioned within four to five cells of the vulva to ensure consistency in size, as cells in BWM taper in width toward the head and tail. For results shown in Fig,1C, D, F, and G, four muscle cells in ten animals of each genotype and age were analyzed, in each of two independent replicates. For results shown in 2B-D and 3A-F, two to four muscle cells in 20 animals of each genotype were analyzed in each of three independent replicates. BWM and muscle cell widths were measured and striations per muscle cell were counted on phalloidin-stained worms (Figs 1C,D, 2B-D, 3A-F), as previously (Sundaramurthy et al., 2020). In young larvae, it is difficult to unambiguously identify BWM striations based on F-actin stain. Therefore, ATN-1::mCH-positive dense bodies were used to identify striations for counting in BWM cells across multiple ages (Fig.1F). Non-parallel striations, identified as those that intersect other striations, were also counted in these cells (Fig.1G). The prevalence of non-parallel striations was also quantified by acquiring single focal plane images of MYO-3 immunostain in six wild type and 13 *fhod-1(ΔFH2)* worms, counting all non-parallel striations in all intact muscle cells, and measuring the combined length of all visible MYO-3 striations in those cells. The prevalence of non-parallel striations was then expressed as the number of occurrences per 10,000 µm of striation measured in each image (Fig.1I). ATN-1::mCH-positive dense bodies were also used to identify striations for determining the effects of drug treatments on muscle cell growth (Fig.S3).

FFT analyses on ATN-1::mCH fluorescence intensity profiles (Table 1, Fig.S2) were performed as we have previously for ATN-1 immunostain (Sundaramurthy et al., 2020). Fluorescence profiles were measured over eight dense body-containing striations in one muscle cell (or all striations in two muscle cells for L1 larvae) in each of ten animals per strain and per developmental stage. To quantify the effects of 24 h levamisole treatment on dense body morphology (Fig.6C), confocal images showing four to five muscle cell striations were acquired from 15 ATN-1 immunostained worms per genotype per condition, in each of three independent replicates. Images were deconvolved as above, and maximum intensity projections were processed through a macro that set a binary threshold of grey values 20-255 to generously mask ATN-1 signal with minimal background, and analyzed particles as % Area. This assay was performed three times. All quantitative measures were performed while blinded to strain genotype.

### FRAP analysis

For analysis of dense body protein dynamics in live animals (Fig.4C,D), L4 stage larvae were immobilized in 3.5 µl polystyrene nanobead suspension (2.5% by volume, 0.1 µm diameter; Polysciences, Warrington, PA) sandwiched between a coverslip and a 10% agarose pad. Worms were viewed for PAT-3::GFP or ATN-1::mCH fluorescence on an SP8 Laser Scanning Confocal Microscope, as above. For each of four (for PAT-3::GFP FRAP) or five (for ATN-1::mCH FRAP) worms of each genotype tested, four regions of interest (ROIs), each encompassing one dense body, were photobleached twice under the FRAP application with 100% laser power, and subsequently imaged for 40 min (for PAT-3::GFP) or 72 min (for ATN-1::mCH). Animals were confirmed to be alive at the end of each experiment by observation of pharyngeal movements. Fluorescence analysis was conducted in ImageJ, measuring at each time point (t) maximum fluorescence intensity values in same-sized ROIs surrounding the four experimental photobleached dense bodies (F_exp_) and two control unbleached dense bodies (F̄_unb_), as well as mean fluorescence intensity values for same-sized ROIs surrounding three background areas of BWM with no dense bodies or other PAT-3::GFP/ATN-1::mCH-containing structures (F_back_). Percent fluorescence was calculated as:

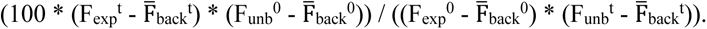

### Pluronic gel burrowing assay

Worms at one day of adulthood were analyzed for the ability to burrow through 26% (w/v) pluronic F-127 (Sigma-Aldrich, St. Louis, MO) gel (Fig.2E), as previously described (Lesanpezeshki et al., 2019). Every 10 min, worms that had burrowed successfully through 0.76 cm gel to OP50-1 chemoattractant on the surface were counted. This assay was performed four times on 30 worms per genotype per replicate.

### Muscle strength measurements using the Nemaflex platform

Strength measurements were conducted using the NemaFlex platform (Fig.1J) as previously described (Rahman et al., 2018). Briefly, at least 30 animals were loaded individually into polydimethylsiloxane microfluidic chambers filled with M9 buffer, where they crawled between free-end deflectable micropillars with diameter, gap, and height of 44 µm, 71 µm, and 87 µm, respectively. Animals were imaged for 1 min at 5 fps at 20 ± 1°C using a Ti-E microscope (Nikon) with an Andor Zyla sCMOS 5.5 camera (Oxford Instruments, Abingdon, United Kingdom). Movies were analyzed using a custom-built image processing software (MATLAB, R2016a; https://github.com/VanapalliLabs/NemaFlex). Muscle strength was calculated based on the maximum pillar deflection identified in each frame, using the Timoshenko theory for an elastic rod (Timoshenko & Gere, 1972). The maximum exerted force *f*_95_ was calculated as the 95^th^ percentile of all maximal deflections for each animal and reported as muscle strength by averaging over the population tested.

### Statistical analysis

Data are expressed as average ± standard deviation, except Fig.1J reporting average ± SEM. Graphs were made in Excel:Windows (version 21H1; Microsoft Corporation, Redmond, WA) or Prism 10 (version 10.1.1; GraphPad Software, Boston, MA). For muscle strength measurements, statistical analysis was performed using a Wilcoxon rank-sum test. For other experiments, where two groups were compared, data were analyzed using a student t-test, and where three or more groups were compared, data were analyzed using a one-way analysis of variation, followed by Tukey’s multiple comparisons *post hoc* test. *p* ≤ 0.05 was considered statistically significant.

## ACKNOWLEDGMENTS

Some worm strains were provided by the CGC, which is funded by NIH Office of Research Infrastructure Programs (P40 OD010440). We thank Peter Calvert for help with FFT analysis, and WormBase and WormBook. R.S.L. would like to thank Anna Lee McKinney for assistance in the generation of RSL62. This work was supported by the National Aeronautics and Space Administration (NNX15AL16G to S.A.V.), University of South Alabama Research Development Award (to R.S.L.), a National Science Foundation EPSCoR Research Infrastructure Improvement Track 4 Grant (Award #1738564 to R.S.L.), the National Institute for Arthritis, Musculoskeletal and Skin Diseases (1R01AR064760 to D.P.), and the Francis Hendricks Endowment Fund (to D.P.).

## SUPPLEMENTARY FIGURE LEGENDS

**Figure S1. Three-dimensional reconstructions from deconvolved confocal z-stacks show BWM from PAT-3::GFP/ATN-1::mCH-expressing animals.** Dense bodies at the ends of striations frequently contain PAT-3::GFP only (*cyan arrowheads*), while dense bodies away from the striation ends have ATN-1::mCH-rich projections (*white arrows*). PAT-3::GFP is also present in M-lines (*white arrowheads*). Dense bodies in *fhod-1(ΔFH2)* animals are short, wide, and sometimes lobulated (*cyan arrows*) compared to wild type. Scale bars, 1 µm.

**Figure S2. Loss of FHOD-1 disrupts dense body spacing in all ages of animals.** Maximum intensity projections of deconvolved confocal z-stacks show ATN-1::mCH-expressing larvae and adults. Scale bars, 4 µm. Dense body spacing along striations was analyzed by performing FFT analysis on ATN-1::mCH fluorescence intensity profiles from images such as these (n = 10 animals per strain per developmental stage per trial, 8 dense body-containing striations from 1 muscle cell or 2 muscle cells for L1 larvae, data shown for two trials, *left* and *right*).

**Figure S3. Assembly of new striations in BWM is inhibited by prolonged treatment with low doses of levamisole or muscimol.** Striations were counted in age-matched ATN-1::mCH-expressing animals that had been grown 24 h in the absence of drug (*vehicle*), or the presence of 2.5 mM levamisole or 10 mM muscimol (n = 10 animals per strain per trial, one muscle cell per animal, two independent trials). Both drugs inhibited addition of new striations in wild type and *fhod-1(ΔFH2)* animals, with levamisole having a much greater effect. Results shown as individual measures and averages ± standard deviation. (**) *p* < 0.01; (***) *p* < 0.001.

## REFERENCES

1. Antoku S, Wu W, Joseph LC, Morrow JP, Worman HJ, Gundersen GG (2019). ERK1/2 phosphorylation of FHOD connects signaling and nuclear positioning alternations in cardiac laminopathy. Dev Cell 51, 602–616.

2. Arribere JA, Bell RT, Fu BXH, Artiles KL, Hartman PS, Fire AZ (2014). Efficient marker-free recovery of custom genetic modifications with CRISPR/Cas9 in *Caenorhabditis elegans*. Genetics 193, 837--846.

3. Ausubel FM, Brent R, Kingston RE, Moore DD, Seidman JG, Smith JA, Struhl K (1992). Short Protocols in Molecular Biology 2nd Edition, New York: Willey.

4. Aydin F, Courtemanche N, Pollard TD, Voth GA (2018). Gating mechanisms during actin filament elongation by formins. Elife 7, e37342.

5. Barstead RJ, Waterston RH (1989). The basal component of the nematode dense-body in vinculin. J Biol Chem 264, 10177–10185.

6. Bremer KV, Wu C, Patel AA, He KL, Grunfeld AM, Chanfreau GF, Quinlan ME (2024). Formin tails act as a switch, inhibiting or enhancing processive actin elongation.

7. Brenner S (1974). The genetics of *Caenorhabditis elegans*, Genetics 77, 71–94.

8. Brouilly N, Lecroisey C, Martin E, Pierson L, Mariol M-C, Qadota H, Labouesse M, Streichenberger N, Mounier N, Gieseler K (2015). Ultra-structural time-course study in the *C. elegans* model for Duchenne muscular dystrophy highlights a crucial role for sarcomere-anchoring structures and sarcolemma integrity in the earliest steps of the muscle degeneration process. Hum Mol Genet 24, 6428–6445.

9. Courtemanche N (2018). Mechanisms of formin-mediated actin assembly and dynamics. Biophys Rev 10, 1553–1569.

10. Dickinson DJ, Ward JD, Reiner DJ, Goldstein B (2013) Engineering the *Caenorhabditis elegans* genome using Cas9-triggered homologous recombination. Nat Methods 10, 1028–1034.

11. Fenix AM, Neininger AC, Taneja N, Hyde K, Visetsouk MR, Garde RJ, Liu B, Nixon BR, Manalo AE, Becker JR, et al. (2018). Muscle-specific stress fibers give rise to sarcomeres in cardiomyocytes. Elife 7, e42144.

12. Finney M, Ruvkun G (1990). The *unc-86* gene product couples cell lineage and cell identity in *C. elegans*. Cell 63, 895–905.

13. Francis GR, Waterston RH (1985). Muscle organization in *Caenorhabditis elegans*: localization of proteins implicated in thin filament attachment and I-band organization. J Cell Biol 101, 1532–1549.

14. Fujimoto N, Kan-O M, Ushimima T, Kage Y, Tominaga R, Sumimoto H, Takeya R (2016). Transgenic expression of the formin protein Fhod3 selectively in the embryonic heart: role of actin-binding activity of Fhod3 and its sarcomeric localization during myofibrillogenesis. PLoS One 11, e0148472.

15. Gettner SN, Kenyon C, Reichardt LF (1995). Characterization of bpat-3 heterodimers, a family of essential integrin receptors in *C. elegans*. J Cell Biol 129, 1127–1141.

16. Gieseler K, Qadota H, Benian GM (2017). Development, structure, and maintenance of *C. elegans* body wall muscle. In: Wormbook, ed. The C. elegans Research Community, doi/10.1895/wormbook.1.81.

17. Gould CJ, Maiti S, Michelot A, Graziano BR, Blanchoin L, Goode BL (2011). The formin DAD domain plays dual roles in autoinhibition and actin nucleation. Curr Biol 21, 384–390.

18. Iskratsch T, Lange S, Dwyer J, Kho AL, dos Remedios C, Ehler E (2010). Formin follows function: a muscle-specific isoform of FHOD3 is regulated by CK2 phosphorylation and promotes myofibril maintenance. J Cell Biol 191, 1159–1172.

19. Henderson HA, Gomez CG, Novak SM, Mi-Mi L, Gregorio CC (2017). Overview of the muscle cytoskeleton. Compr Pysiol 7, 891–944.

20. Huxley H, Hanson J (1954). Changes in the cross-striations of muscle during contraction and stretch and their interpretation. Nature 173, 973–976.

21. Kan-O M, Takeya R, Abe T, Kitajima N, Nishida M, Tominaga R, Kurose H, Sumimoto H (2012). Mammalian formin Fhod3 plays an essential role in cardiogenesis by organizing myofibrillogenesis. Biol Open 1, 889–896.

22. Kooij V, Viswanathan MC, Lee DI, Rainer PP, Schmidt W, Kronert WA, Harding SE, Kass DA, Bernstein SI, Van Eyk JE, et al. (2016). Profilin modulates sarcomeric organization and mediates cardiomyocyte hypertrophy. Cardiovasc Res 110, 238–248.

23. Kovar DR, Kuhn JR, Tichy AL, Pollard TD (2003). The fission yeast cytokinesis formin Cdc12p is a barbed end actin filament capping protein gated by profilin. J Cell Biol 161, 875–887.

24. Kovar DR, Pollard TD (2004). Insertional assembly of actin filament barbed ends in association with formins produces piconewton forces. Proc Natl Acad Sci U S A 101, 14725–14730.

25. Maufront J, Guichard B, Cao-L-Y, Di Cicco A, Jégou A, Romet-Lemonne G, Bertin A (2023). Direct observation of the conformational states of formin mDia1 at actin filament barbed ends and along the filament. Mol Biol Cell 34, ar2.

26. Miller DM, III, Ortiz I, Berliner GC, Epstein HF (1983). Differential localization of two myosins within nematode thick filaments. Cell 34, 477–490.

27. Moerman DG, Williams BD (2006). Sarcomere assembly in *C. elegans* muscle. In: Wormbook, ed. The C. elegans Research Community, doi/10.1895/wormbook.1.81.

28. Moulder GL, Cremona GH, Juerr J, Stirman JN, Fields SD, Martin W, Qadota H, Benian GM, Lu H, Barstead RJ (2010). a-actinin is required for the proper assembly of Z-disk/focal-adhesion-like structures and for efficient locomotion in *Caenorhabditis elegans*. J Mol Biol 403, 516–528.

29. Lesanpezeshki L, Hewitt JE, Laranjeiro R, Antebi A, Driscoll M, Szewczyk NJ, Blawzdziewicz J, Lacerda CMR, Vanapalli SA (2019). Pluronic gel-based burrowing assay for rapid assessment of neuromuscular health in *C. elegans*. Sci Rep 9, 15246.

30. Lesanpezeshki L, Qadota H, Darabad MN, Kashyap K, Lacerda CMR, Szewczyk NJ, Benian GM, Vanapalli SA (2021). Investigating the correlation of muscle function tests and sarcomere organization in *C. elegans*. Skelet Muscle 11, 20.

31. Mi-Mi L, Votra S, Kemphues K, Bretscher A, Pruyne D (2012). Z-line formins promote contractile lattice growth and maintenance in striated muscles of *C. elegans*. J Cell Biol 198, 87–102.

32. Mi-Mi L, Pruyne D (2015). Loss of sarcomere-associated formins disrupts Z-line organization, but does not prevent thin filament assembly in *Caenorhabditis elegans* muscle. J Cytol Histol 6, 318.

33. Moseley JB, Sagot I, Manning AL, Xu Y, Eck MJ, Pellman D, Goode BL (2004). A conserved mechanism of Bni1- and mDia1-induced actin assembly and dual regulation of Bni1 by Bud6 and profilin. Mol Biol Cell 15, 896–907.

34. Moss BM, Refsnes PE, Abildgaard A, Nicolaysen K, Jensen J (1997). Effects of maximal effort strength training with different loads on dynamic strength, cross-sectional area, load-power and load-velocity relationships. Eur J Appl Physiol 75, 193–199.

35. Nishida E, Iida K, Yonezawa N, Koyasu S, Yahara I, Sakai H (1987). Cofilin is a component of intranuclear and cytoplasmic actin rods induced in cultured cells. Proc Natl Acad Sci U S A 84, 5262–5266.

36. Ochoa JP, Sabater-Molina M, García-Pinilla JM, Morgensen J, Restrepo-Córdoba A, Palomino-Doza J, Villacorta E, Martinez-Moreno M, Ramos-Maqueda J, Zorio E, et al. (2018) Formin homology 2 domain containing 3 (FHOD3) is a genetic basis for hypertrophic cardiomyopathy. J Am Coll Cardiol 72, 2457–2467.

37. Ono K, Ono S (2002). Tropomyosin inhibits ADF/cofilin-dependent actin filament dynamics. J Cell Biol 156, 1065–1076.

38. Ono K, Ono S (2009). Actin-ADF/cofilin rod formation in *Caenorhabditis elegans* muscle requires a putative F-actin binding site of ADF/cofilin at the C-terminus. Cell Motil Cytoskeleton 66, 398–408.

39. Ono S, Abe H, Obinata T (1996). Stimulus-dependent disorganization of actin filaments induced by overexpression of cofilin in C2 myoblasts. Cell Struct Funct 21, 491–499.

40. Ono S, Baillie DL, Benian GM (1999). UNC-60B, an ADF/cofilin family protein, is required for proper assembly of actin into myofibrils in *Caenorhabditis elegans* body wall muscle. J Cell Biol 145, 491–502.

41. Paix A, Folkmann A, Rasoloson D, Seydoux G (2015). High efficiency, homology-directed genome editing in *Caenorhabditis elegans* using CRISPR-Cas9 ribonucleoprotein complexes. Genetics 201, 47–54.

42. Patel AA, Oztug Durer ZA, van Loon AP, Bremer KV, Quinlan ME (2018). *Drosophila* and human FHOD family formin proteins nucleate actin filaments. J Biol Chem 293, 532–540.

43. Pimm ML, Hotaling J, Henty-Ridilla JL (2020). Profilin choreographs actin and microtubules in cells and cancer. Int Rev Cell Mol Biol 355, 155–204.

44. Plenefisch JD, Zhu X, Hedgecock EM (2000). Fragile skeletal muscle attachments in dystrophic mutants of *Caenorhabditis elegans*: isolation and characterization of the *mua* genes. Development 127, 1197–1207.

45. Polet D, Lambrechts A, Ono K, Mah A, Peelman F, Vandekerckhove J, Baillie DL, Ampe C, Ono S (2006). *Caenorhabditis elegans* expresses three functional profilins in a tissue-specific manner. Cell Motil Cytoskeleton 63, 14–28.

46. Pruyne D, Evangelista M, Yang C, Bi E, Zigmond S, Bretscher A, Boone C (2002). Role for formins in actin assembly: nucleation and barbed-end association. Science 297, 612–615.

47. Rahman M, Hewitt JE, Van-Bussel F, Edwards H, Blawzdziewicz J, Szewczyk NJ, Driscoll M, Vanapalli SA (2018). NemaFlex: a microfluidics-based technology for standardized measurements of muscular strength of *C. elegans*. Lab Chip 18, 2187–2201.

48. Ramabhadran V, Gurel PX, Higgs HN (2012). Mutations to the formin homology 2 domain of INF2 protein have unexpected effects on actin polymerization and severing. J Biol Chem 287, 34234–34245.

49. Refai O, Smit RB, Votra S, Pruyne D, Mains PE (2018). Tissue-specific functions of *fem-2*/PP2c phosphatase and *fhod-1*/formin during *Caenorhabditis elegans* embryonic morphogenesis. G3 (Bethesda) 8, 2277–2290.

50. Romero S, Le Clainche C, Didry D, Egile C, Pantaloni D, Carlier M-F (2004). Formin is a processive motor that requires profilin to accelerate actin assembly and associated ATP hydrolysis. Cell 119, 419–429.

51. Sagot I, Rodal AA, Moseley J, Goode BL, Pellman D (2002). An actin nucleation mechanism mediated by Bni1 and profilin. Nat Cell Biol 4, 626–631.

52. Schneider CA, Rasband WS, Eliceri KW (2012). NIH image to ImageJ: 25 years of image analysis. Nature Methods 9, 671–675.

53. Shwartz A, Dhanyasi N, Schejter ED, Shilo B-Z (2016). The *Drosophila* formin Fhos is a primary mediator of sarcomeric thin-filament array assembly. Elife 5, e16540.

54. Sundaramurthy S, Votra S, Laszlo A, Davies T, Pruyne D (2020). FHOD-1 is the only formin in *Caenorhabditis elegans* that promotes striated muscle growth and Z-line organization in a cell autonomous manner. Cytoskeleton (Hoboken) 77, 422–441.

55. Taniguchi K, Takeya R, Suetsugu S, Kan-O M, Narusawa M, Shoise A, Tominaga R, Sumimoto H (2009). Mammalian formin fhod3 regulates actin assembly and sarcomere organization in striated muscles. J Biol Chem 284, 29873–29881.

56. Timoshenko S, Gere JM (1971). Mechanics of Materials, New York: Van Nostrand Reinhold Company.

57. Ushijima T, Fujimoto N, Matsuyama S, Kan-O M, Kiyonari H, Shioi G, Kage Y, Yamasaki S, Takeya R, Sumimoto H (2018). The actin-organizing formin protein Fhod3 is required for postnatal development and functional maintenance of the adult heart in mice. J Biol Chem 293, 148–162.

58. Vizcarra CL, Bor B, Quinlan ME (2014). The role of formin tails in actin nucleation, processive elongation, and filament bundling. J Biol Chem 289, 30602–30613.

59. Xu Y, Moseley JB, Sagot I, Poy F, Pellman D, Goode BL, Eck MJ (2004). Crystal structures of a formin homology-2 domain reveal a tethered dimer architecture. Cell 116, 711–723.

60. Waterston RH (1988). Muscle. In: The Nematode Caenorhabditis elegans, ed. W. B. Wood, New York: Cold Spring Harbor Laboratory Press, 281–335.

61. Wooten EC, Hebl VB, Wolf MJ, Greytak SR, Orr NM, Draper I, Calvino JE, Kapur NK, Maron MS, Kullo IJ, et al. (2013) Formin homology 2 domain containing 3 variants associated with hypertrophic cardiomyopathy. Circ Cardiovasc Genet 6, 10–18.

62. Yingling CV, Pruyne D (2021). FHOD formin and SRF promote post-embryonic striated muscle growth through separate pathways in *C. elegans*. Exp Cell Res 398, 112388.

63. Zeiser E, Frøkjær-Jensen C, Jorgensen E, Ahringer J (2011). MosSCI and gateway compatible plasmid toolkit for constitutive and inducible expression of transgenes in the *C. elegans* germline. PLoS One 6, e20082.

64. Zhou Q, Wei S-S, Wang H, Wang Q, Li W, Li G, Hou J-W, Chen X-M, Chen J, Xu W-P, et al. (2017). Crucial role of ROCK2-mediated phosphorylation and upregulation of FHOD3 in the pathogenesis of angiotensin II-induced cardiac hypertrophy. Hypertension 69, 1070–1083.

