## Supplemental Methods for "FHOD-1/profilin-mediated actin assembly protects sarcomeres against contraction-induced deformation in *C. elegans*"

### *atn-1* genomic editing

The *C. elegans* operon CEOP5346 located on chromosome V consists of the genes *atn-1* (W04D2.1; 5632 nt) followed by *srt-39* (W04D2.2; 1184 nt). The 837 nt intergenic region includes the *atn-1* 3' UTR (405 nt). The presence of a *srt-39* 5' UTR within the intergenic region is not documented but is likely limited in size due to the presence of a potential start codon 19 nt upstream of the actual start codon. The most likely 3' splice site (TCCTGAG|A) that ends 2 nt upstream is a poor fit to the consensus, while a slightly better conforming site (TTTTTAG|C) ends 26 nt upstream from the actual start codon. The use of two alternate start sites and an exon cassette (for exon 4) produces four Atn-1 isoforms, that are all composed from the last five constitutive exons at the 3' end of the gene (exons 5 – 9). The Cas9 target sequence (GTGACGTAATCGAATGATCG) used for the genomic insertion of the BiTS cassette is located on the non-coding strand 25 – 45 nt upstream of the *atn-1* stop codon and the PAM (TGG) is located 46 – 48 nt upstream of the *atn-1* stop codon. Five silent mutations adjacent to the PAM site were included in the repair plasmid to remove the Cas9 target sequence (CGA-TCA -> AGG-AGC, Arg-Ser). No change could be made in the PAM site itself without altering the identity of the amino acid encoded (CCA, Pro). The mCherry-ICR-GFP sequence was inserted prior to the *atn-1* stop codon. This resulted in an in-frame fusion of atn-1, followed by a Gly5-Ala linker, and mCherry, ending the Y37E3.8 3' UTR. After trans-splicing, the second gene encoded GFP, beginning with the *rla-1* 5'UTR and ending with the *atn-1* 3'UTR, and the *srt-39* gene became the third gene in the operon.
