## Supplementary figures and images for "FHOD-1/profilin-mediated actin assembly protects sarcomeres against contraction-induced deformation in *C. elegans*"

### Fig.S1

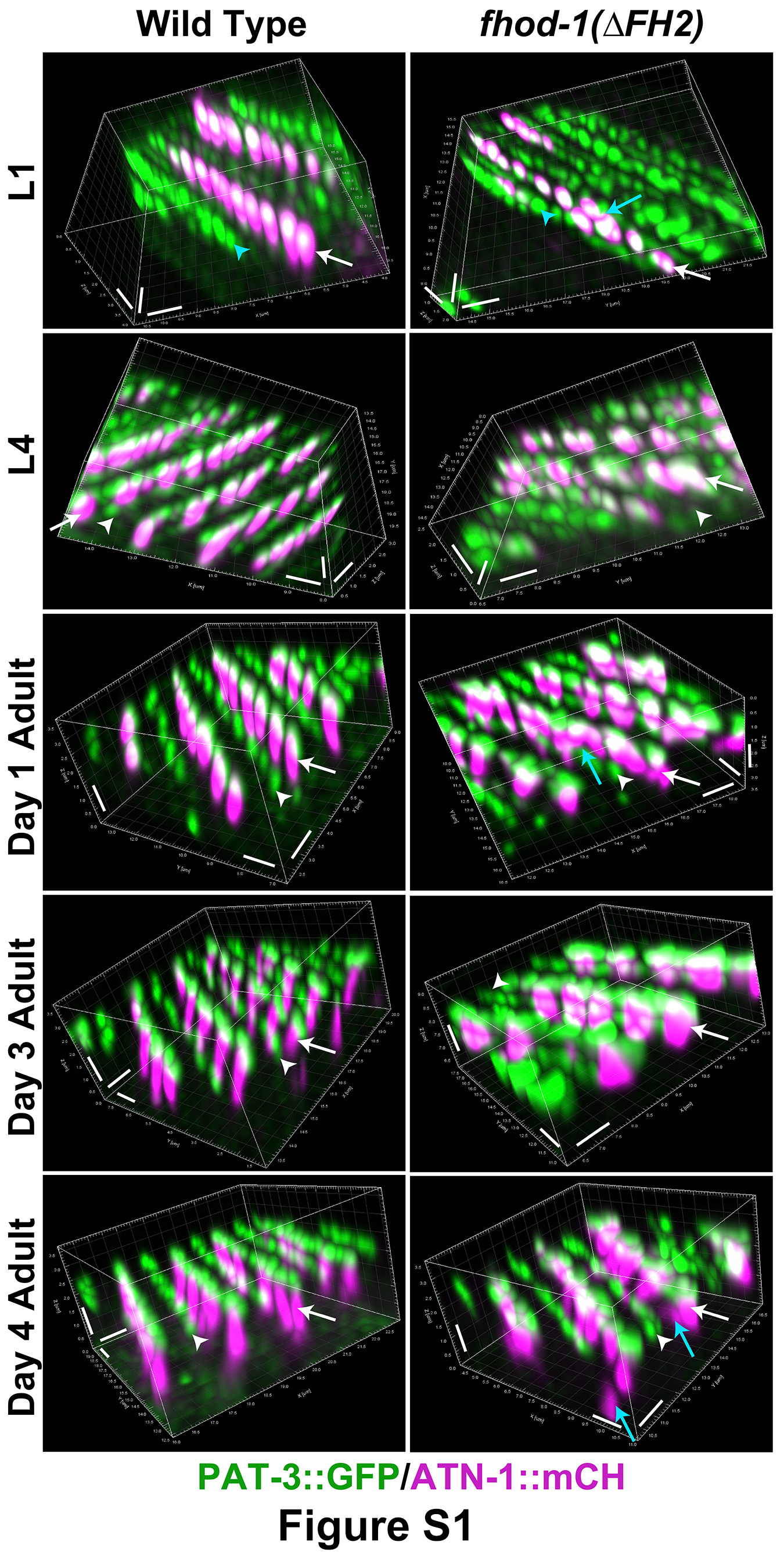

### Fig.S2

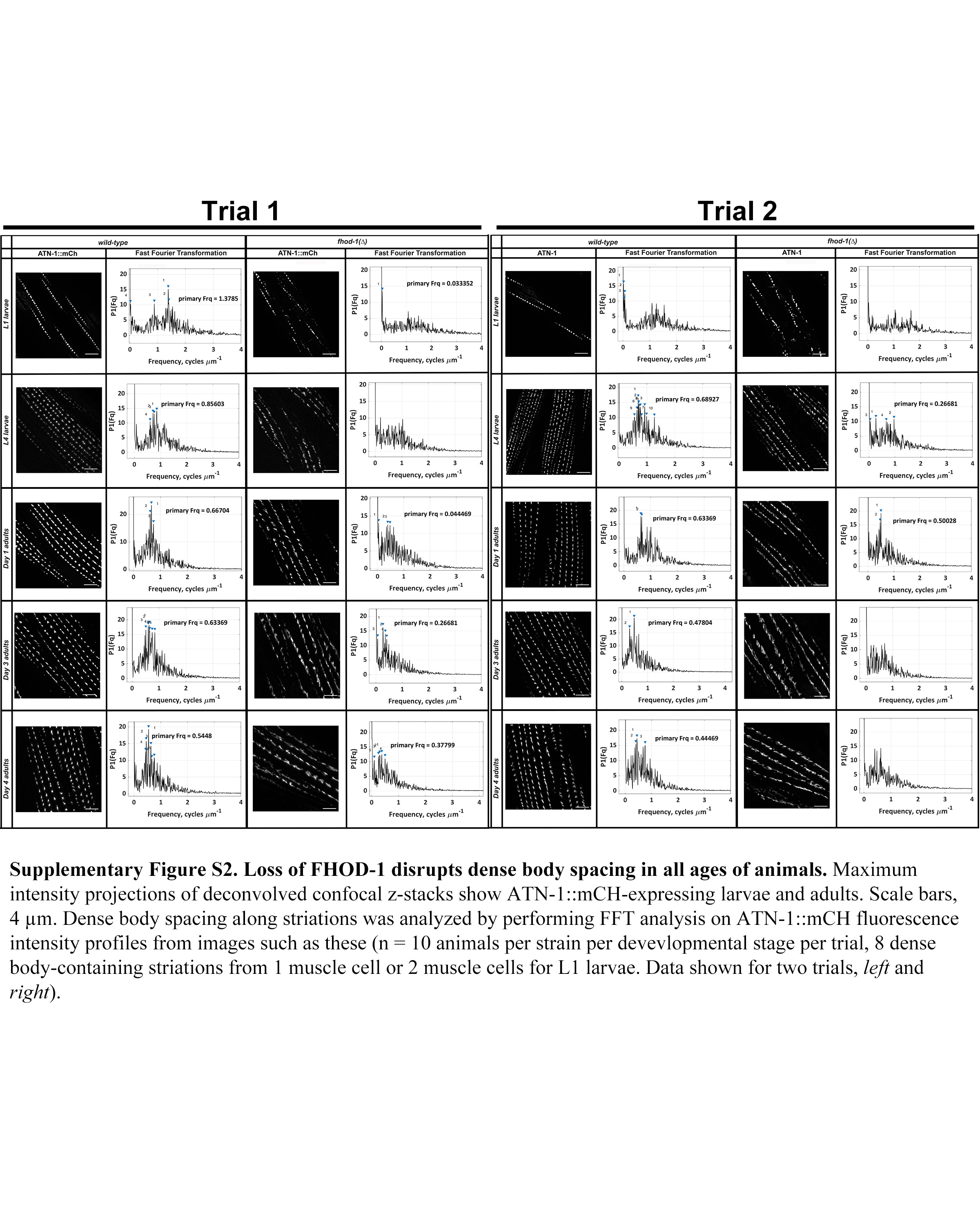

### Fig.S3

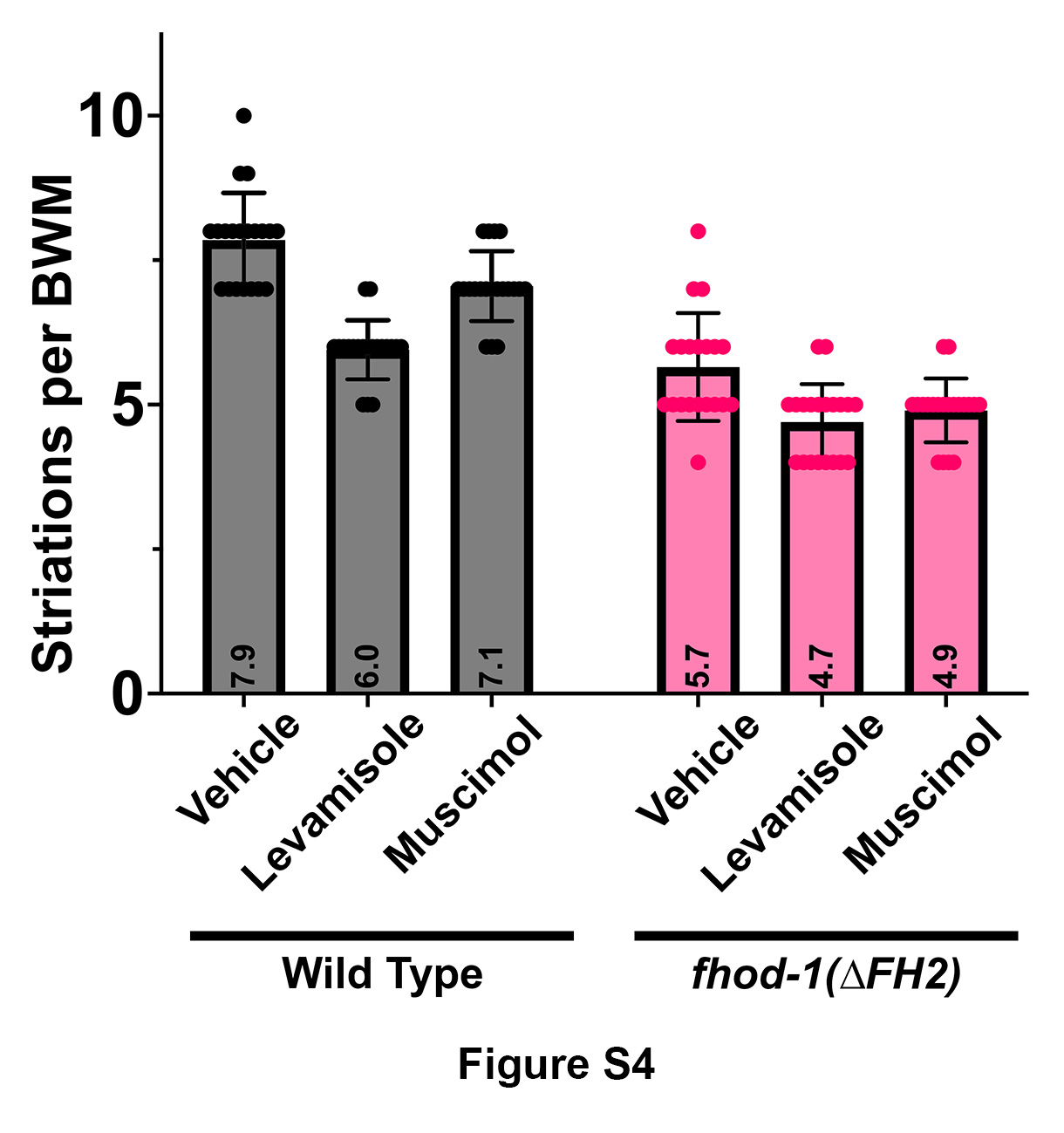
